# Polysaccharides from *Balanophora harlandii* Hook: Isolation, Characterization, and Anti-Inflammation Activities

**DOI:** 10.1101/2023.09.27.559774

**Authors:** Yuanyang Li, Xueqing Li, Qi Yuan, Leiqi Zhu, Fangqi Xia, Yaqi Wang, Mengzhen Xue, Yumin He, Chengfu Yuan

## Abstract

*Balanophora harlandii* Hook (*B. harlandii)*, a folk medicine, has been traditionally employed to treat traumatic bleeding, gastroenteritis, icteric hepatitis, hemorrhoids, and other conditions. In this work, polysaccharides with anti-inflammatory effects were extracted and purified from *B. harlandii.* The extraction conditions were optimized, and the properties of one purified neutral fraction, denoted as BHPs-W-S3, were analyzed. Gel permeation chromatography (GPC) was carried out to measure the molecular weight. The structure of BHPs-W-S3 was assessed based on monosaccharide composition analysis, Fourier transform infrared (FT-IR) spectroscopy, methylation analysis, and nuclear magnetic resonance (NMR) spectroscopy. BHPs-W-S3 has a molecular weight of 14.1 kDa, and its three main monosaccharides are glucose, galactose, and mannose with a molar ratio of 6.4:1.7:1.1. Its main chain consists of →6)-α-D-Glcp-(1→, →4,6)-α-D-Glcp-(1→, →6)-β-D-Galp-(1→, →3,6)-β-D-Galp-(1→, and it has branch chains at the O-4 and/or O-3 positions. In addition, *in vitro* experiments show that the polysaccharides from *B. harlandi* can decrease the phosphorylation level of p65 and IKB-α in LPS-induced RAW264.7 cells to reduce the expression of the pro-inflammatory genes such as TNF-α, IL-6, and IL-1β.

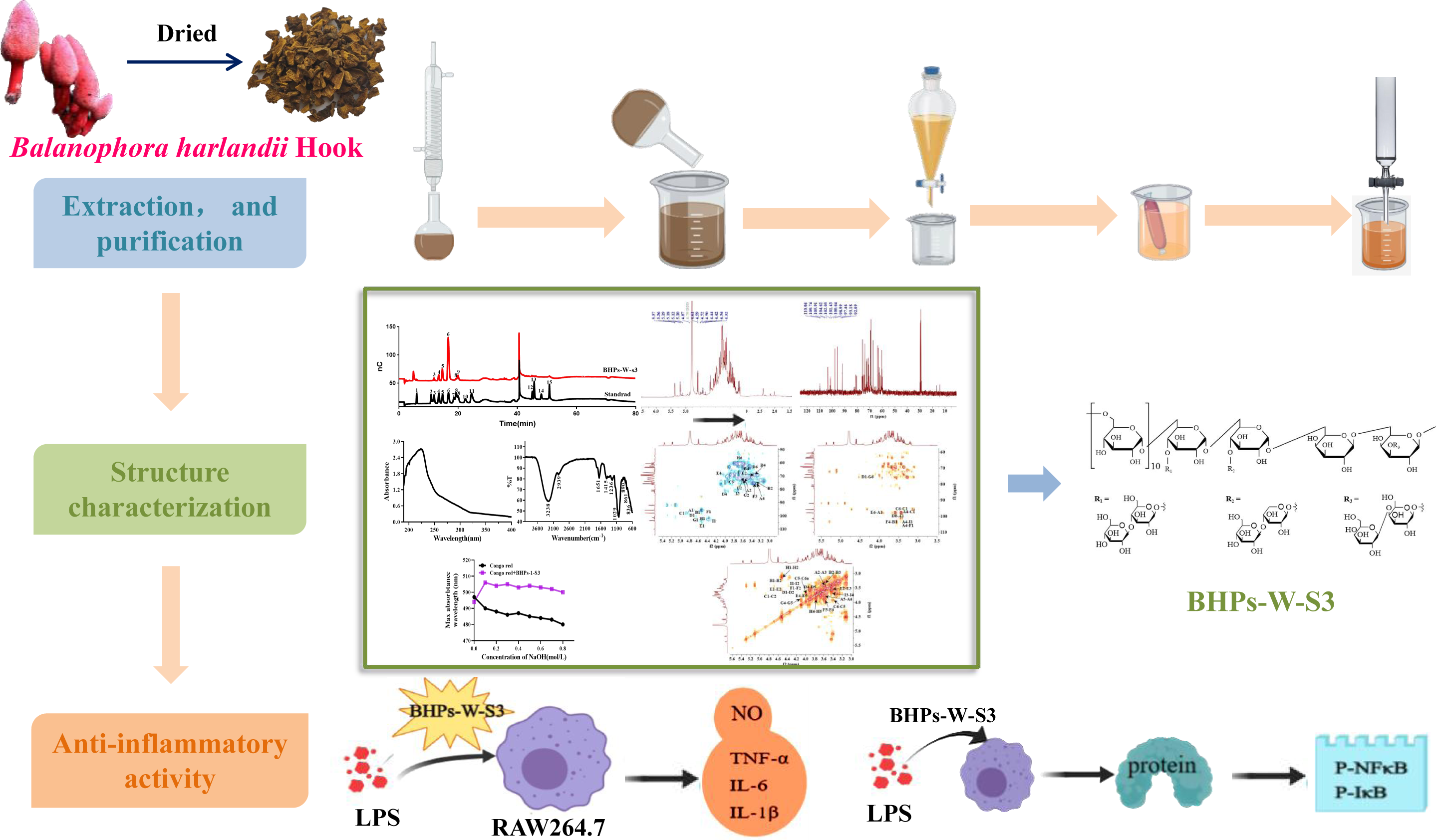

## 1. Introduction

Polysaccharides from plants have shown diverse biological activities and health benefits. They are typically non-toxic and rarely have severe side effects (Xu et al., 2009). They have been used as adjuvants in treating various diseases, as they can function as an antioxidant (Kerboua et al., 2021), fight infection, reduce inflammation, enhance immunity (Liu et al., 2020; Wang et al., 2021), moderate blood sugar, and prevent cancer (Jia et al., 2021).

*Balanophora harlandii* Hook (*B. harlandii)*, also known as Wenwang Yizhibi, is a perennial herb of the Balanophoraceae family. In accordance with traditional Chinese herbal texts, the entire herb of *B. harlandii* possesses a calming nature, a slightly astringent and bitter taste, and is reputed for its efficacy in clearing heat and detoxifying, nourishing yin and blood, transforming blood stasis, and promoting tissue regeneration (Herbals, 1999). It is perceived as famed and precious traditional medicine in local area of Tujia, Mao and other ethinc minotities in China, which was commonly applied to the treatment of wounds, gastrointestinal bleeding, gonorrhea, syphilis, etc (Yuanbei et al., 2017). Some chemical components have been isolated from *B. harlandii* and studied, including tannins, flavonoids, terpenoids, sterols, and polysaccharides (Wang et al., 2012; Li et al., 2021), wherein *Balanophora* polysaccharide was considered to be one of the main effective substances of *B. harlandii*.

Our preliminary research shows that the *Balanophora* polysaccharides (BPs) have outstanding anti-inflammatory effects. In vitro colitis animals treated with dextran sulfate sodium and in *vivo* RAW 264.7 cells stimulated by lipopolysaccharide (LPS) both showed reduced expression of pro-inflammatory genes when BPs was present (Guo et al., 2020). The BPs also restrains the proliferation of ovarian cancer cells by modulating the P53 pathway (Qu et al., 2020). Further research is needed to clarify the physicochemical properties, structural characteristics, and biological functions of BPs.

In this work, we developed a protocol to extract crude polysaccharides from the whole plant of *B. harlandii* by hot water, then separated the polysaccharide using a DE-52 column, and subsequently purified it with a Sephadex G-100 column to obtain a water-soluble neutral polysaccharide (BHPs-W-S3). The physicochemical properties of BHPs-W-S3 were analyzed by gel permeation chromatography (GPC), Fourier transform infrared (FT-IR) spectroscopy, gas chromatography-mass spectrometry (GC-MS) and nuclear magnetic resonance (NMR) spectroscopy. The anti-inflammatory effects of BHPs-W-S3 were evaluated with RAW 264.7 cells using standard biochemical assays. The findings contribute to the informed utilization of *Balanophora* plants as a significance economic resource in medicinal applications.

## 2. Materials and Methods

### 2.1. Materials

#### 2.1.1. Isolation and purification of polysaccharides from *B. harlandii*

The samples of *B. harlandii* were obtainde from Wufeng County, Hubei, China and identified by Prof. Y-B Wang at the Biotechnology Research Center of China Three Gorges University. The voucher specimens were stored in the specimen room at the Institute of Metabolic Diseases Drug Research of China Three Gorges University.

Fig. 1 illustrates the workflow to isolate and purify the BHPs. The baked-dried *B. harlandii* (420 g) were pulverized into powder and sieved with 60 mesh, followed by 95% ethanol (2L) solution degreasing for 4 h. The resultant filter residues was dried by a vacuum-drying oven at 55 °C by a vacuum-drying oven, and boiled thrice with water in a ratio of 1:20 (w/v). The combined aqueous extract (20 L) was centrifuged and concentrated to about 20% of the initial volume at 55 °C under reduced pressure, and 95% ethanol (12 L) was then added. The precipitates were collected by centrifugation, washed twice with 95% ethanol, dissolved in water, and finally lyophilized to give the crude BHPs.

**Fig.1.**
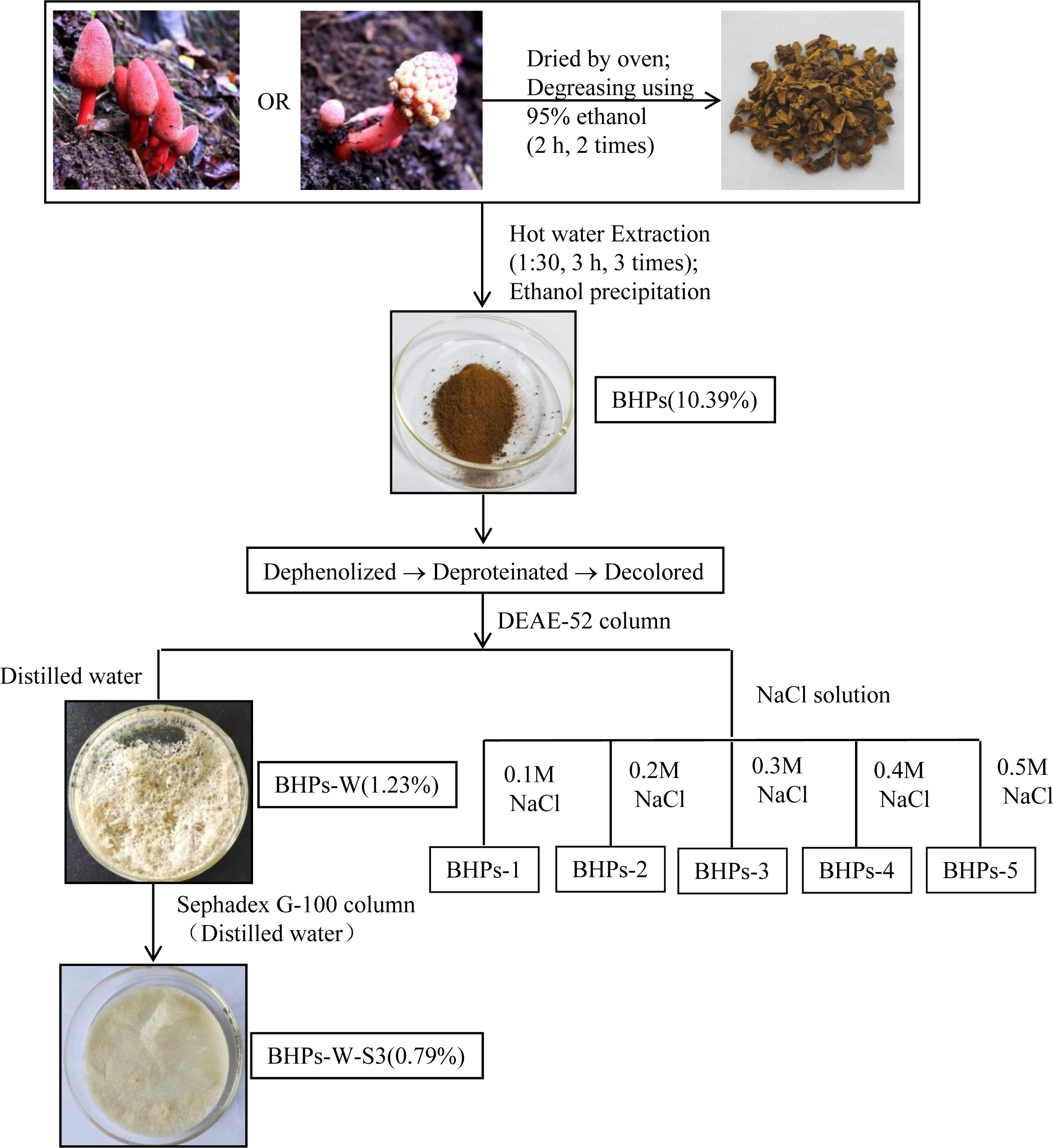
The extraction and isolation of polysaccharides from *Balanophora harlandii* Hook.

Crude BHPs were dissolved in water and liquid-liquid fractionation with ethyl acetate (EtoAc) on a separating funnel to remove phenolic. The lower aqueous layer was combined with 4 M CaCl_2_ (1:1 v/v), boiled for 1 min to remove proteins (Zeng et al., 2019). Subsequently, the supernatant collected by centrifugation was treated with 1% ammonium oxalate under magnetic stirring for 30 min, and after a second centrifugation, the supernatant was decolored by heating at 70 °C for 30 min with activated carbon. The aqueous solutions was filtered after cooling to room temperature, and the filtrate was then dialyzed (Mw cutoff at 3500 Da) for 48 h in ultra-pure water. The purified BHPs (denoted as BHPs-PE) were obtained after the dialysate was concentrated under reduced pressure and lyophilized.

For further purification, BHPs-PE (0.5 g) was loaded on a DE-52 anion exchange column (2.4 × 70 cm) and eluted with a step-wise gradient (distilled water to 0.5 M aqueous NaCl) at 1 mL/min (Fig. 1). The eluate was monitored by phenol-sulfuric acid method (DuBois et al., 1956). The polysaccharide fraction corresponding to the elution with distilled water, denoted as BHPs-W, was loaded on a Sephadex G-100 column (2.4 × 120 cm) and eluted with distilled water at 0.1 mL/min. The fractions accordant to the prominent maxima was combined, dialyzed, and lyophilized to give BHPs-W-S3, which was stored for further analysis.

#### 2.1.2. Single-factor and orthogonal design optimization of crude polysaccharide extraction

Three single factors were firstly examined to optimize the extraction, and the extraction yield corresponding to each extraction condition was calculated. Specifically, the ratio of water-to-raw material, the duration of extraction, and the repetition of extractions were varied from 10:1 to 50:1 mL/g, 1 to 4 h, and 1 to 5 times, respectively. The total polysaccharide was determined by the phenol-sulfuric acid method (DuBois et al., 1956). The equation presented below was utilized to compute the extract yield of BHPs:

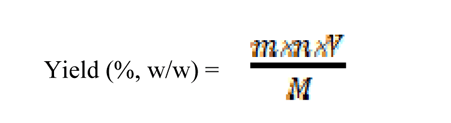

where m (mg) is glucose content, *n* is the dilution coefficient, *V* is the volume of the test solution, and *M* (g) is the dry weight of *B. harlandii*.

Subsequent to the outcomes of the single-factor analysis, three-factors-three-levels of an orthogonal design was carried out to further explore the optimum extraction process in terms of the yield of total polysaccharide. Table 1 displays the L_9_(3^3^) orthogonal table.

**Table 1.** Independent variables Levels for Orthogonal Tests.

| Level | Factor |  |  |
| --- | --- | --- | --- |
|  | (A) Ratio of water-to-raw material (mL/g) | (B) Extraction time (h) | (C) The number of extraction (times) |
| 1 | 20:1 | 2 | 1 |
| 2 | 30:1 | 3 | 2 |
| 3 | 40:1 | 4 | 3 |

Table 1 Independent variables Levels for Orthogonal Tests.
| Level | Independent variables |  |  |
| --- | --- | --- | --- |
|  | (A) Ratio of water to raw material (mL/g) | (B)Extraction time (h) | (C)The number of Extraction (times) |
| 1 | 20:1 | 2 | 1 |
| 2 | 30:1 | 3 | 2 |
| 3 | 40:1 | 4 | 3 |

### 2.2. Characterizations

#### 2.2.1. Chemical components

The total carbohydrate content was determined by the phenol-sulfuric acid method using glucose as the standard (DuBois et al., 1956). The proteins content was analyzed by the Coomassie brilliant blue method using bovine serum albumin (BSA) as the standard (Bradford, 1976). The total phenol content was analyzed by the Folin–Ciocalteu method using gallic acid as the standard (Singleton and Rossi, 1965).

#### 2.2.2. Molecular weight

The molecular weight of BHPs-W-S3 was measured by gel permeation chromatography using a tandem gel column (8 × 300 mm) and a RI-10 differential detector. The column was loaded with an aliquot (20 μL) of BHPs-W-S3 prepared by dissolving 5 mg BHPs-W-S3 in 1 mL water, and then eluted with 0.05 M aqueous NaCl at 0.6 mL/min. The column was maintained at 40 °C. The calibration curves were constructed using dextran standards (Fig. S2) and the retention time was plotted against the logarithm of the molecular weight. The molecular weight of BHP-W-S3 was inferred from the curves.

#### 2.2.3. Monosaccharide composition

The sample of BHPs-W-S3 (10 mg) was hydrolyzed in 2 M trifluoroacetic acid (TFA, 10 mL) at 120 °C for 4 h. The supernatant was collected after centrifugation at 4000 rpm for 5 min and blown to dryness with a stream of nitrogen to give the hydrolysate of BHPs-W-S3, which was then dissolved in distilled water (5 mL) and analyzed by ion chromatography on a Dionex Carbopac PA20 column (3 × 150 mm) by gradient elution with a binary system of Solution A (H_2_O) and Solution B (15 mM NaOH and 100 mM NaOAc) at a flow rate of 0.3 mL/min.

#### 2.2.4. Ultraviolet and FT-IR spectroscopic analysis

To measure the UV absorption, BHPs-W-S3 (10 mg) was dissolved in distilled water (5 mL) and the absorption over 190–400 nm was measured on a UV-Vis spectrophotometer using a quartz cell with 1 cm path length.

Dried BHPs-W-S3 (*ca*. 2 mg) was pulverized with KBr powder. Subsequently, the powder was blended and compressed, and examined by FT-IR spectroscopy within the wavelength range of 600-4000 cm^-1^ to identify their characteristic peaks.

#### 2.2.5. Methylation analysis

The methylation of BHPs-W-S3 was analyzed according to a published procedure (Luo et al., 2018). Specifically, BHPs-W-S3 (3 mg) was dissolved in anhydrous DMSO (1 mL) and methylated using NaH and CH_3_I. The methylated product was hydrolyzed by TFA, reduced by NaBH_4_, and acetylated by acetic anhydride. Finally, the acetylated material was extracted thrice with CH_2_Cl_2_ for GC-MS analysis on an Agilent setup that includes an RXI-5 SIL MS column.

#### 2.2.6. Nuclear magnetic resonance spectroscopy analysis

The BHPs-W-S3 (30 mg) was dissolved in 1 mL of D_2_O and subjected to lyophilization followed by three rounds of repeated freeze-thaw cycles. Then 0.6 mL D_2_O was added followed by transferring the sample solution to an NMR tube. 1D NMR (^1^H spectrum, ^13^C spectrum) and 2D NMR (COSY spectrum, HSQC spectrum, HMBC spectrum) were analyzed by using a Bruker Avance 600 MHz spectrometer.

#### 2.2.7. Congo red test

To a mixture of the BHPs-W-S3 solution (2 mg/mL, 1 mL) and the Congo red solution (0.2 mM, 1.5 mL) was added 2 M NaOH, and the final concentration of NaOH was adjusted to 0–0.8 mol/L. The mixture was maintained at room temperature for 5 min before the absorption over 400–600 nm was scanned. The pure Congo red solution was used as the control.

### 2.3. Anti-inflammatory activity

#### 2.3.1. Cells and other materials

The Shanghai Academy of Life Sciences’ cell resource center in Shanghai, China, provided the RAW 264.7 cell lines. Primary antibodies against β-actin, IKB-α, p-IKB-α, p65, and p-p65 were procured from Wanlebio, situated in Shenyang, China. Tianjin Chemical Reagent Co. Ltd., Tianjin, China, provided the other chemical reagents utilized.

#### 2.3.2. RAW 264.7 cells culture

The RAW 264.7 cells were cultured in DMEM enriched with 10% heat-inactivated FBS and 1% penicillin/streptomycin at 37 °C under an atmosphere containing 5% CO_2_.

#### 2.3.3. MTT assay

The RAW 264.7 cells were seeded in 96-well plates (1 × 10^5^ cell/mL, 100 μL per well), and phosphate-buffered saline (PBS) was added into the surrounding wells to prevent edge effects. After having reached the logarithmic growth stage, the cells were incubated with crude BHPs or BHPs-W-S3 (25–200 μg/mL) in DMEM containing 1 μg/mL LPS for 24 h. The blank and model group were pure DMEM and 1 μg/mL LPS in DMEM, respectively. The MTT solution (5 mg/mL, 10 μL) was then added to each well, and the cells were cultured at 37 °C for another 4 h in an incubator with 5% CO_2_. Afterwards, the supernatant was removed, DMSO (100 μL) was added, the plates were shaken for 10 min to ensure the complete dissolution of crystals, and the absorbance at 570 nm was measured with a microplate reader (Thermo Electron, TYPE1500-458, USA).

#### 2.3.4. Determination of nitric oxide (NO) production

The NO production of RAW 264.7 cells was determined according to a published protocol (Jang et al., 2022). The RAW 264.7 cells were incubated for 24 h with 1 μg/mL LPS in DMEM in 96-well plates (5 × 10^5^ cells/mL) to simulate inflammation, and then incubated with crude BHPs or BHPs-W-S3 (25‒100 μg) for another 24 h. The positive control group was Dexamethasone (DEX). Afterwards, the supernatant was collected, and the same amount of Griess reagent was added. The NO production was calculated from the absorption at 540 nm measured with a microplate reader.

#### 2.3.5. Real-time PCR

The RAW 264.7 cells’ total RNA was extracted employing the Trizol reagent. Its absorption evaluated from 260 to 280 nm, the isolated RNA can be ascertained the concentration and purity. According to the instructions from the manufacturer, the cDNA was synthesized by means of the PrimeScript RT Reagent Kit. The conditions for RT-qPCR were as follows: pre-incubation at 50 °C for 2 min, denaturation at 95 °C for 10 min, and 40 cycles of 95 °C for 15 s and 60 °C for 60 s. The internal control was GAPDH. The data were analyzed by the 2^−ΔΔCt^ method. Table 2 lists the sequences of the primer used.

**Table 2.**
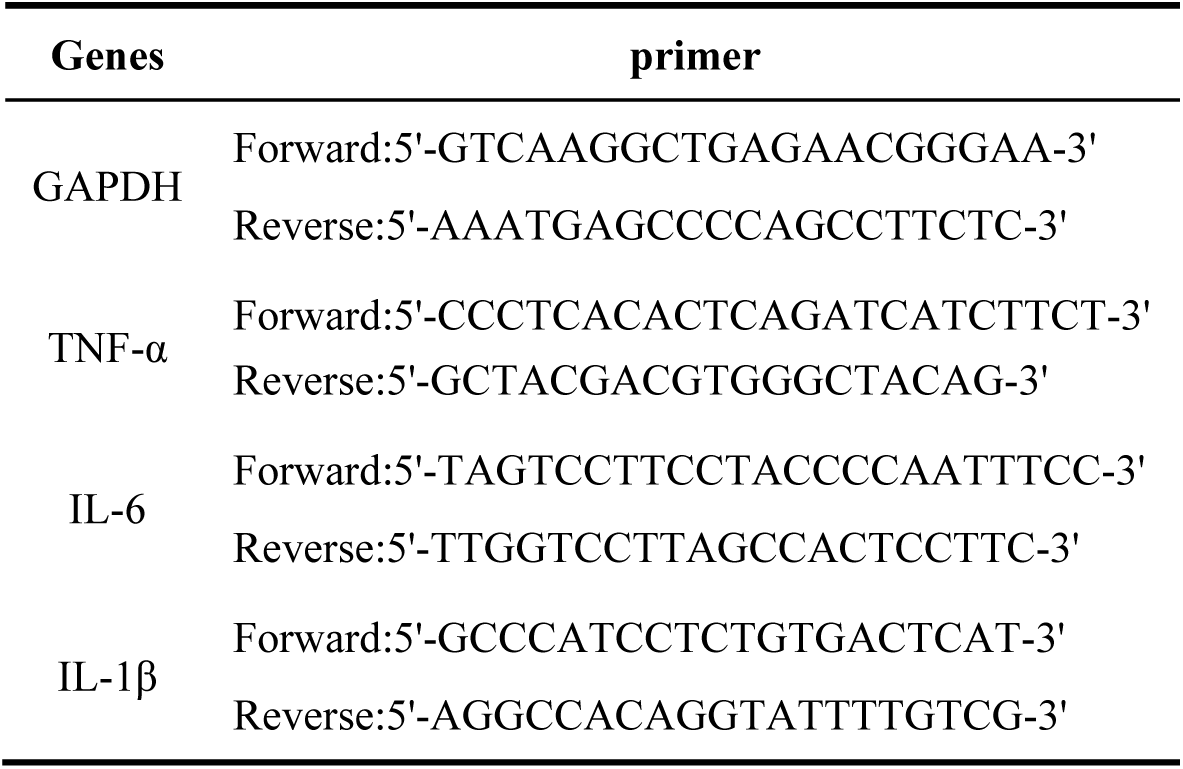
Primers for the required genes.

#### 2.3.6. Western blot analysis

Its outcome of lysing centrifuged at 12000g and 4 °C for 20 min, the RAW 264.7 cells were lysed in RIPA buffer at 0 °C. The concentration of protein was measured by means of the BCA protein assay kit. The proteins (30 μg per lane) were separated by SDS-PAGE and transferred onto polyvinylidene difluoride (PVDF) membranes. The membranes were blocked in a solution of Tris-buffered saline (TBS)-Tween 20 containing 5% skimmed dry milk and incubated with the primary antibody at 4 °C overnight, then visualized by electrochemiluminescence.

### 2.4. Statistical analysis

All measurements were repeated three times. Data are presented as the mean ± standard deviation (SD) and analyzed by one-way ANOVA using SPSS Statistics 21.0. The criterion for statistical significance is *p* < 0.05.

## 3. Results and Discussions

### 3.1. Optimization of the extraction

#### 3.1.1. Single-factor test

The effect of the water-to-raw material ratio (10–50 mL/g) on the extraction yield was firstly studied by running the extraction once for 1 h. The yield of crude BHPs rises sharply when the water use increases from 10 to 20 mL/g but then dwindles when more water is used (Fig. 2A). Increasing the water use improves the solubility of polysaccharides from herbs by enabling the thorough penetration of solvents and diffusion of solutes (Gu et al., 2020). Therefore, water-to-raw material of 20–40 mL/g was chosen for the orthogonal experiment.

**Fig.2.**
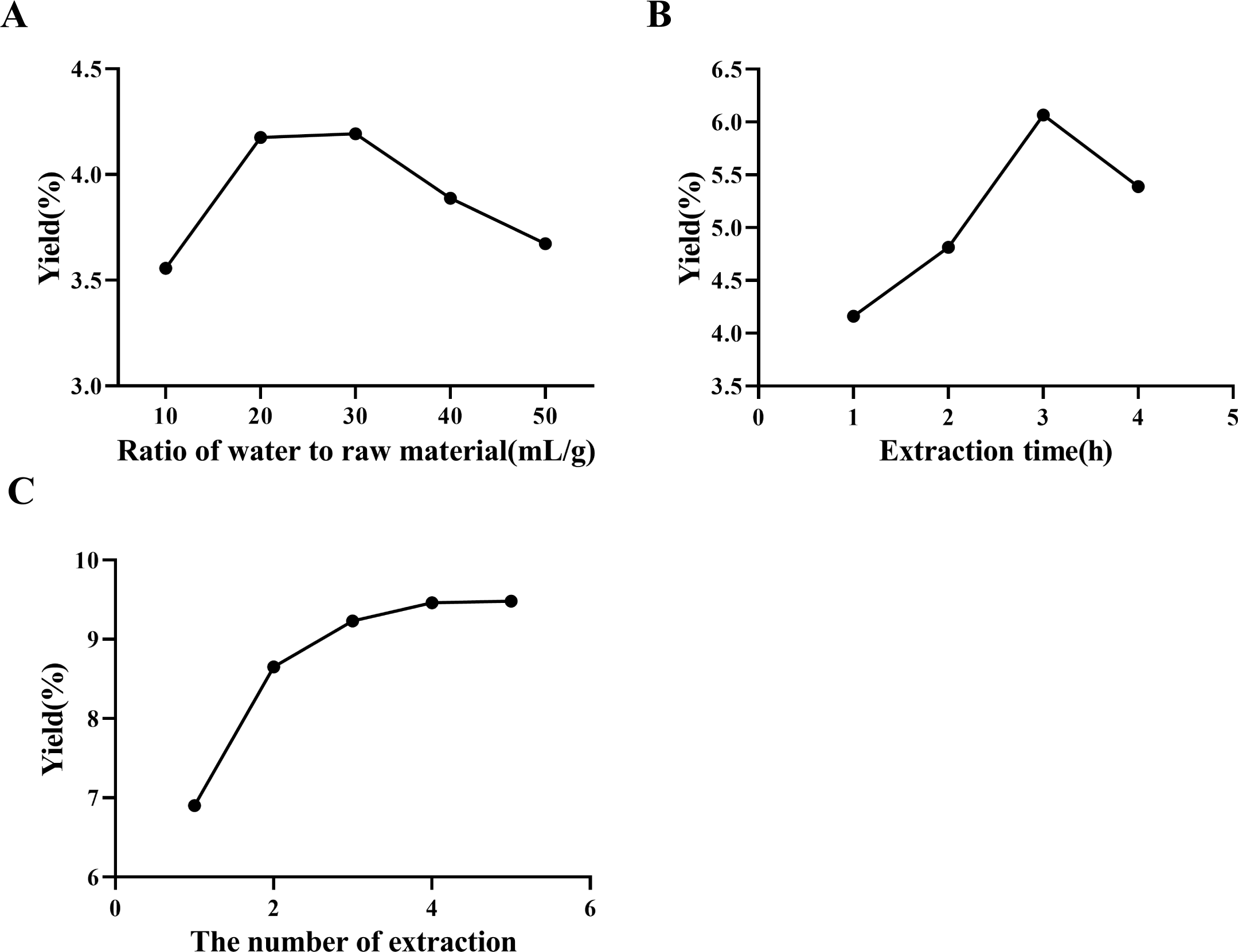
Single factor tests to optimize of the extraction yield. (A) Ratio of water to raw material, (B) duration of extraction, (C) the number of extraction.

Under the fixed conditions of a water-to-raw material ratio of 30 mL/g and a single extraction heating cycle, the effect of varying extraction durations (1, 2, 3, and 4 h) on the yield of BHPs was examined. As depicted in Figure 2B, the yield of BHPs increased rapidly from 1 to 3 h, with a peak of 6.07% at 3 h, and then gradually decreased with further increase in extraction durations. Since a prolonged extraction may cause the breakdown of polysaccharides (Liu and Li, 2020), the extraction time was varied from 2 to 4 h in the subsequent orthogonal experiment.

Finally, the extraction was run 1 to 5 times, each time for 3 h using 30 mL/g water, and the yield of crude BHPs from the combined extract was calculated. Figure 2C shows that the extraction yield increases monotonously with the number of extractions. As the number of extractions increases, the volume of solution consumed also increases, and the corresponding energy consumption rises significantly. In view of the extraction efficiency, the subsequent orthogonal design assessed 1–3 repetitions of extractions.

#### 3.1.2. Orthogonal test results analysis

Three key factors were considered in the orthogonal test to maximize the extraction yield of BHPs: (A) water-to-raw material ratio, (B) extraction time, and (C) number of extraction. Table 3 shows that their contribution to the extraction efficiency falls in the order of C > B > A, and the extraction yield varies considerably among different levels. The optimum extraction condition is as follow: a ratio of water-to-raw material 30 mL/g, trice extractions at 3 h per extraction. To test the reliability of the given method, BHPs were extracted for six batches under the above condition. The RSD value was < 2%, indicating consistency in predictive values.

**Table 3.**
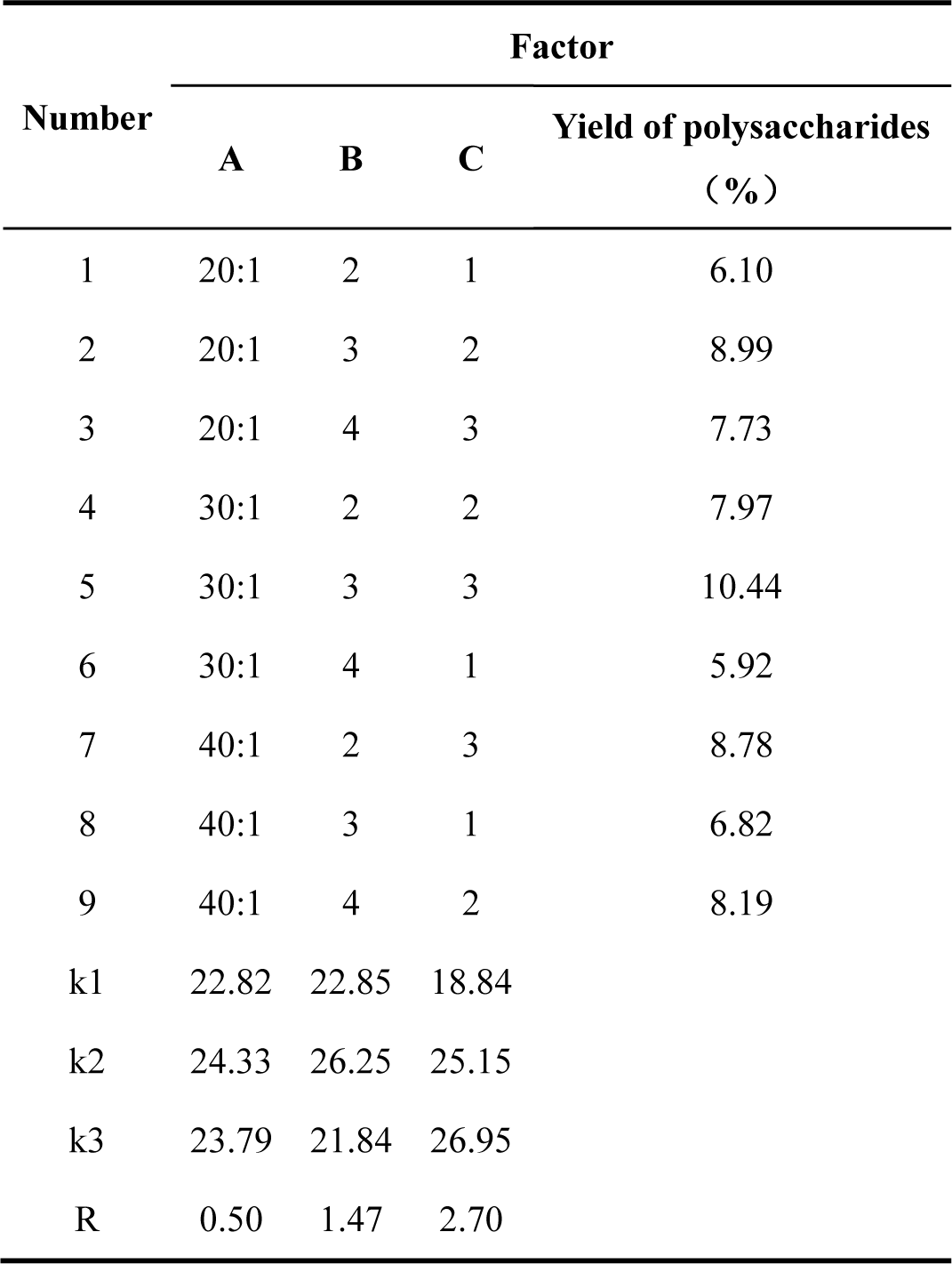
Results of orthogonal experiment.

### 3.2. Physicochemical characteristics of the crude BHPs

#### 3.2.1. Purification

The crude BHPs were obtained in approximately 10.39% mass yield from the dried *B. harlandii* plant after hot water extraction and ethanol precipitation. Separation using a DE-52 cellulose column gave a neutral fraction (Fig. 3A) that was purified further on a Sephadex G-100 gel column (Fig. 3B) to afford BHPs-W-S3 in 0.79% yield. Table 4 shows that the total sugar, protein, and phenol contents of crude BHPs are 35.15%, 9.94%, and 23.25%, respectively, while BHPs-W-S3 has a total sugar content of 82.71%, virtually no protein, and very little phenol.

**Fig.3.**
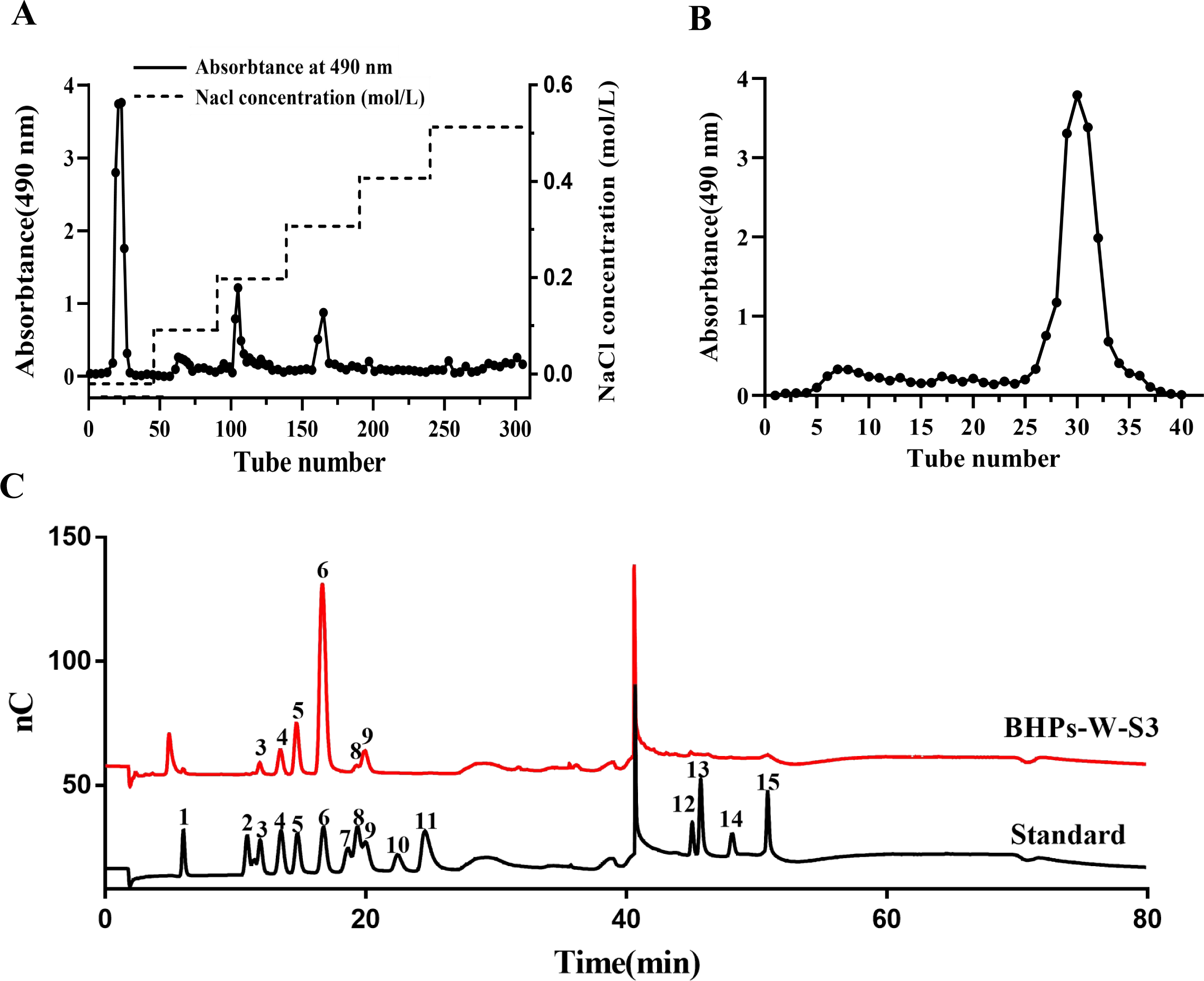
(A) Elution of BHPs-PE on the DE-52 anion exchange column. (B) Elution of the neutral faction on the Sephadex G-100 gel filtration column, (C) Analysis of the monosaccharide composition of BHPs-W-S3 by ion chromatography. 1: Fuc; 2: Rha; 3: Ara; 4: GlcN; 5: Gal; 6: Glc; 7: GlcNC; 8: Xyl; 9: Man; 10: Fru; 11: Rib; 12: GalA; 13: GulA; 14: GlcA; 15: ManA.

**Table 4.** Yield and chemical components of BHPs and BHPs-W-S3.

| Sample | BHPs | BHPs-W-S3 |
| --- | --- | --- |
| Yield (%) | 10.39 | 0.79 |
| Total sugar (%) | 35.15±0.38 | 82.71±0.26 |
| Total protein (%) | 9.94±1.76 | - |
| Total phenolic (%) | 23.25±0.43 | 1.90±0.47 |

-, undetectable

#### 3.2.2. Monosaccharide composition and molecular weight

The M_w_ and M_n_ of BHPs-W-S3 are calculated from the calibration curves (Fig. S3) to be 14.1 and 10.5 kDa, respectively, and the polydispersity index (PI) of BHPs-W-S3 is 1.34. The molar ratios of monosaccharides in BHPs-W-S3 are estimated by comparison with the peaks areas of standard monosaccharides in chromatography (Fig. 3C, Table 5). The black curve in Fig. 3C is the spectrum of the standard monosaccharides. The numbers mark the different monosaccharides, and their molar ratio is calculated according to the formula C(standard)/A(standard) = C(sample)/A(sample).

**Table 5.** The monosaccharide composition and molecular weight of the BHPs-W-S3.

| Sample | Sugar(Mole Ratio) |  |  |  |  |  | Mw(kDa) | Mn(kDa) | PI |
| --- | --- | --- | --- | --- | --- | --- | --- | --- | --- |
|  | Ara | GlcN | Gal | Glc | Xyl | Man |  |  |  |
| BHPs-W-S3 | 0.3 | 0.3 | 1.7 | 6.4 | 0.2 | 1.1 | 14.1 | 10.5 | 1.34 |

BHPs-W-S3 is mainly consisted of glucose, galactose, and mannose, and has minor amounts of arabinose, glucosamine hydrochloride, and xylose. It can be inferred that BHPs-W-S3 has relatively homogeneous heteropolysaccharides that are predominantly composed of glucose, galactose, and mannose monosaccharides.

### 3.3. Structural characterization of BHPs-W-S3

#### 3.3.1. UV-Vis and FT-IR spectroscopy

No absorption peaks are found at either 260 nm or 280 nm in the UV spectra of BHPs-W-S3 (Fig. 4A), which indicates the near absence of nucleic acids or proteins in the analyte. The peak absorption at 230 nm can be attributed to carbohydrates containing aromatic groups, phenols, and certain hydrocarbons.

**Fig.4.**
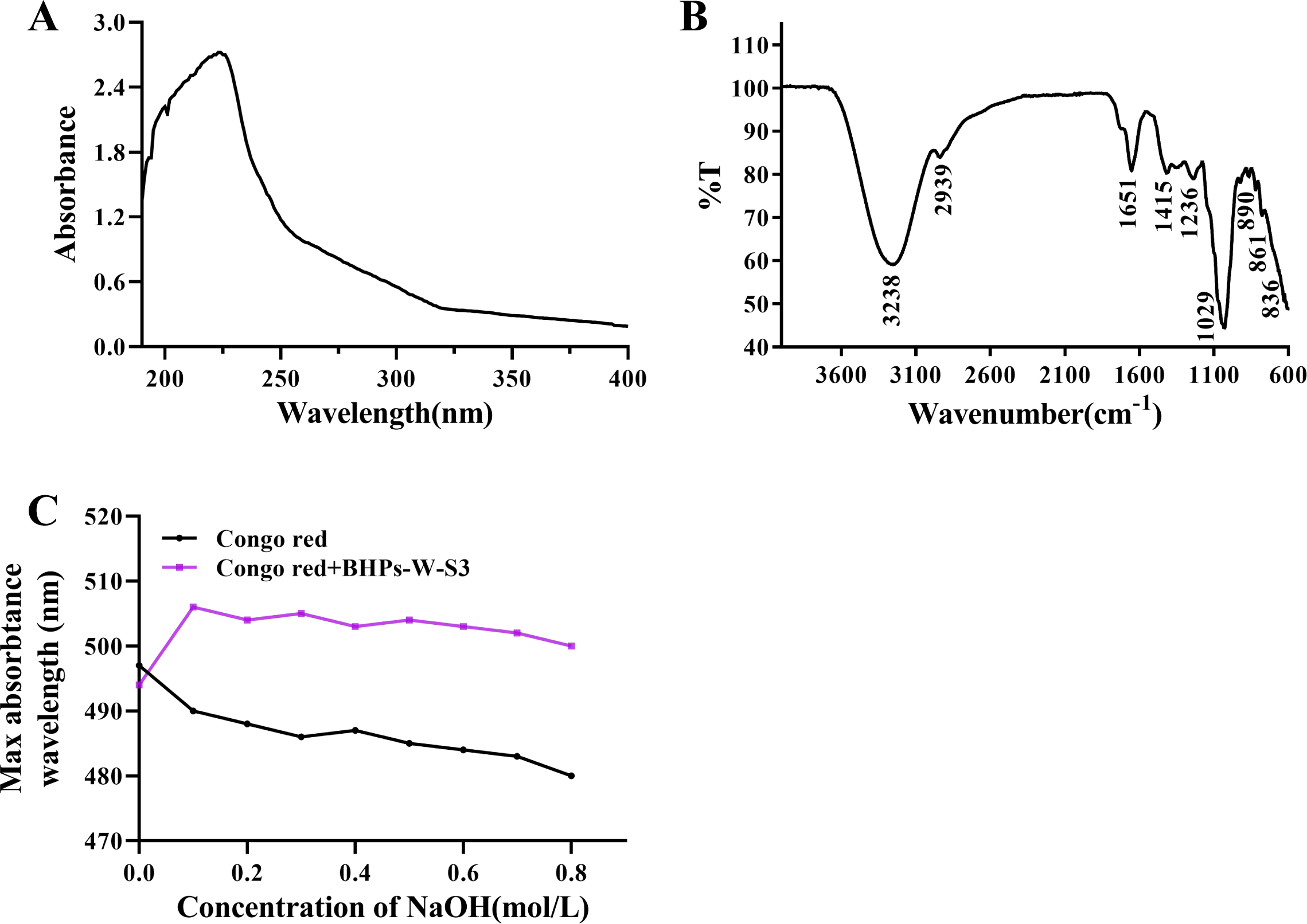
(A) UV-vis spectra and (B) FT-IR spectra of BHPs-W-S3. (C) Maximum absorption wavelength of the Congo red/BHPs-W-S3 mixture with varying concentration of NaOH.

In the FT-IR spectra of BHPs-W-S3 (Fig. 4B), the broad peak at 3238 cm^−1^ can be ascribed to the stretching vibration of ‒OH, and the weak signal at 2939 cm^−1^ arises from the C‒H stretching vibration (Huang and Huang, 2020). The strong absorption at 1029 cm^−1^ can be related to the C‒OH, C‒O‒C, and C‒C stretching vibrations that correspond to the skeleton of the pyran ring structure (Li et al., 2019). The weak absorptions at 890 and 836 cm^−1^ may be related to the β-and α-glycosidic linkages of glucosyl units, respectively (Arab et al., 2021).

#### 3.3.2. GC-MS

Nine methylated and acetylated derivatives (Table 6) can be identified by comparing the major mass fragments in the GC-MS spectra of BHPs-W-S3 (Fig. S1-S2) with the reported data (Li et al., 2019; Ma et al., 2022), i.e., →4)-Xylp-(1→, T-Glcp-(1→, T-Galp-(1→, →4)-Galp-(1→, →4)-Glcp-(1→, →6-Glcp-(1→, →6)-Galp-(1→, →4,6)-Glcp-(1→, and →3,6)-Galp-(1→. Some Glc and Gal residues in BHPs-W-S3 appear to be terminal residues. Thus, T-Glcp and/or T-Galp residues are probably coupled to the 1,4-linked Glcp, 1,4-linked Galp, 1,4,6-linked Glcp, and 1,3,6-linked Galp in the O-4/O-3 position.

**Table 6.** Methylation analysis of BHPs-W-S3.

| **Methylated sugar** | **Mass fragments (m/z)** | **Molar ratio** | **Type of linkage** |
| --- | --- | --- | --- |
| 2,3-Me<sub>2</sub>-Xylp | 43,71,87,99,101,117,129,161,189 | 0.41 | →4)-Xylp-(1→ |
| 2,3,4,6-Me<sub>4</sub>-GlcP | 43,71,87,101,117,129,145,161,205 | 4.28 | GlcP-(1→ |
| 2,3,4,6-Me<sub>4</sub>-Galp | 43,71,87,101,117,129,145,161,205 | 0.75 | Galp-(1→ |
| 2,3,6-Me<sub>3</sub>-Galp | 43,87,99,101,113,117,129,131,161,173,233 | 0.84 | →4)-Galp-(1→ |
| 2,3,6-Me<sub>3</sub>-GlcP | 43,87,99,101,113,117,129,131,161,173,233 | 0.56 | →4)-GlcP-(1→ |
| 2,3,4-Me<sub>3</sub>-GlcP | 43,87,99,101,117,129,161,189,233 | 1.08 | →6)-GlcP-(1→ |
| 2,3,4-Me<sub>3</sub>-Galp | 43,87,99,101,117,129,161,189,233 | 1.20 | →6)-Galp-(1→ |
| 2,3-Me<sub>2</sub>-GlcP | 43,71,85,87,99,101,117,127,159,161,201,261 | 0.68 | →4,6)-GlcP-(1→ |
| 2,4-Me<sub>2</sub>-Galp | 43,87,117,129,159,189,233 | 0.20 | →3,6)-Galp-(1→ |

#### 3.3.3. NMR

The NMR spectroscopy allows the evaluation of anomeric configurations, sugar unit sequences, linkage patterns, number and relative content of the sugar residues, etc. (Hu et al., 2015; Zhang et al., 2017). In general, 5.0 ppm is the threshold value for the proton signal to distinguish between pyranose conformations; H-1 chemical shifts > 5.0 ppm are in α-configuration, otherwise they are in β-configuration; and the 3.5‒4.0 ppm region reflects the H2‒H6 on the sugar ring (Kim et al., 2022). The ^1^H NMR spectrum of BHPs-W-S3 (Fig. 5A) has various signals in 4.5‒5.5 ppm, which suggests the presence of both α-and β-configurations. The overlapping peaks in 3.0‒4.0 ppm can be attributed to the H2‒H6 of heteropolysaccharides. The region of 90‒120 ppm in the ^13^C NMR spectra (Fig. 5B) corresponds to the anomeric carbon, further confirming that BHPs-W-S3 is a heteropolysaccharide. These conclusions are consistent with the FT-IR analysis’s findings. At the same time, DEPT was carried out to identify the existence of primary, secondary and tertiary carbon atoms. In DEPT-135 spectra Fig. 5C, the secondary carbon peaks at 67.8, 62.1, 67.3, 63.8, 70.2, 61.7, 67.8, 62.5, 64.8, 65.3 ppm suggested the glucopyranosyl substitution at C-6.

**Fig.5.**
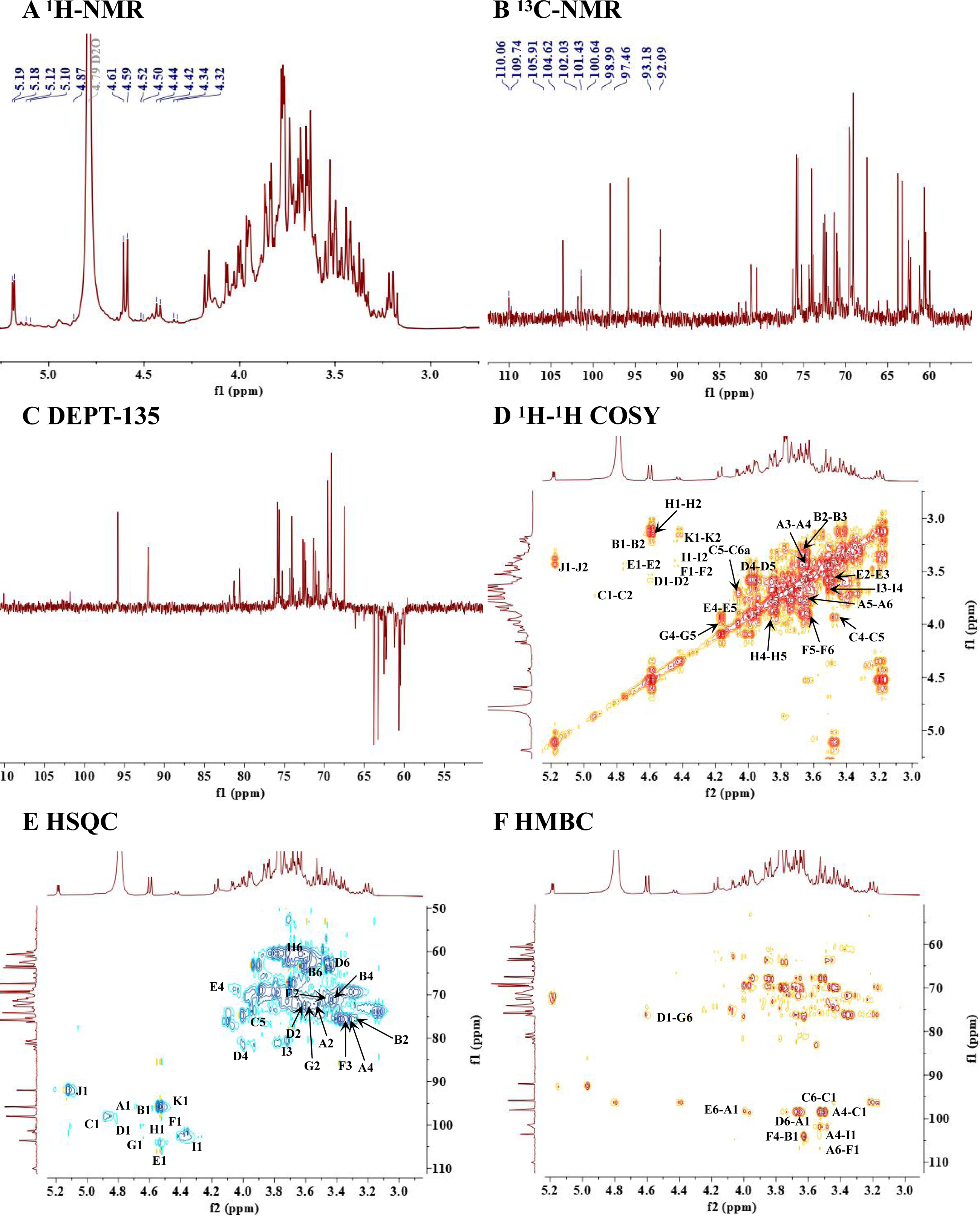
NMR analysis of BHPs-W-S3. (A) ^1^H spectrum, (B) ^13^C spectrum, (C) DEPT-135 spectrum, (D) ^1^H-^1^H COSY spectrum, (E) HSQC spectrum, (F) HMBC spectrum.

Table 7 compiles the assignment of ^1^H and ^13^C NMR spectra. Eleven residues are identified by combining the ^1^H-^1^H COSY (Fig. 5D) and HSQC (Fig. 5E) spectrum of BHPs-W-S3, which are →4,6)-α-D-Glcp-(1→, β-D-Glcp-(1→, →6)-α-D-Glcp-(1→, →4)-β-D-Galp-(1→, →6)-β-D-Galp-(1→, →4)-β-D-Glcp-(1→, →3,6)-β-D-Galp-(1→, β-D-Galp-(1→, →4)-β-D-Xylp-(1→, →6)-α-D-Gal and →6)-β-D-Gal. The proton/carbon chemical shifts at the anomeric position of the glycosyl residues in BHPs-W-S3 are 4.87/99.6, 4.52/104.2, 4.86/99.4, 4.51/105.5, 4.44/108.8, 4.33/104.2, 4.55/104.3, 4.45/104.5, 4.36/104.1, 5.09/93.6 and 4.51/97.4 ppm respectively. Other monosaccharides are not detectable because they make up a relatively small percentage and are susceptible to interference in methylation and NMR analysis.

**Table 7.** ^1^H and ^13^C NMR chemical shifts of BHPs-W-S3 recorded in D_2_O.

| Glycosyl residues | H1/C1 | H2/C2 | H3/C3 | H4/C4 | H5/C5 | H6a/C6 | H6b |
| --- | --- | --- | --- | --- | --- | --- | --- |
| $\rightarrow 4,6)\text{-}\alpha\text{-D-Glcp-(1}\rightarrow$ | 4.87 | 3.49 | 3.67 | 3.36 | 3.57 | 3.85 | 3.67 |
| A | 99.62 | 72.86 | 74.02 | 77.1 | 70.69 | 67.78 |  |
| $\beta\text{-D-Glcp-(1}\rightarrow$ | 4.52 | 3.29 | 3.61 | 3.34 | 3.44 | 3.66 | 3.83 |
| B | 104.2 | 74.17 | 75.25 | 71.15 | 74.25 | 62.12 |  |
| $\rightarrow 6)\text{-}\alpha\text{-D-Glcp-(1}\rightarrow$ | 4.86 | 3.53 | 3.67 | 3.47 | 3.86 | 3.57 | 3.71 |
| C | 99.40 | 72.73 | 74.63 | 70.87 | 71.47 | 67.27 |  |
| $\rightarrow 4)\text{-}\beta\text{-D-Galp-(1}\rightarrow$ | 4.51 | 3.6 | 3.69 | 4.00 | 3.63 | 3.43 | 3.57 |
| D | 105.47 | 73.21 | 73.11 | 82.02 | 76.01 | 63.79 |  |
| $\rightarrow 6)\text{-}\beta\text{-D-Galp-(1}\rightarrow$ | 4.44 | 3.46 | 3.62 | 4.05 | 3.87 | 3.89 | 3.83 |
| E | 108.8 | 72.02 | 74 | 69.82 | 74.81 | 70.2 |  |
| $\rightarrow 4)\text{-}\beta\text{-D-Glcp-(1}\rightarrow$ | 4.33 | 3.3 | 3.44 | 3.67 | 3.47 | 3.95 | 3.57 |
| F | 104.16 | 74.17 | 76.72 | 82.04 | 76.69 | 61.67 |  |
| $\rightarrow 3,6)\text{-}\beta\text{-D-Galp-(1}\rightarrow$ | 4.55 | 3.62 | 3.81 | 4.08 | 3.87 | 3.85 | 3.45 |
| G | 104.3 | 72.79 | 77.98 | 70.05 | 73.34 | 67.78 |  |
| $\beta\text{-D-Galp-(1}\rightarrow$ | 4.45 | 3.27 | 3.45 | 3.66 | 3.8 | 3.64 | 3.77 |
| H | 104.48 | 74.66 | 76.62 | 77.28 | 75.01 | 62.52 |  |
| $\rightarrow 4)\text{-}\beta\text{-D-Xylp-(1}\rightarrow$ | 4.36 | 3.33 | 3.5 | 3.71 | 3.42 | 3.57 | |
| I | 104.07 | 76.65 | 73.69 | 81.22 | 65.23 |  |  |
| $\rightarrow 6)\text{-}\alpha\text{-D-Gal}$ | 5.09 | 3.41 | 3.61 | 3.31 | 4.06 | 3.93 | 3.58 |
| J | 93.62 | 72.52 | 73.82 | 77.2 | 75.71 | 64.76 |  |
| $\rightarrow 6)\text{-}\beta\text{-D-Gal}$ | 4.51 | 3.17 | 3.34 | 3.61 | 3.97 | 3.57 | 3.42 |
| K | 97.43 | 75.21 | 76.84 | 74.08 | 75.91 | 65.33 |  |

The structural skeleton of carbohydrates can be identified based on the HMBC spectrum by correlating remote carbons and protons (Ren et al., 2019). The split cross peaks of C1/H6 at 99.4/3.85 ppm in the HMBC spectrum (Fig. 5F) reveal that →6)-α-D-Glcp-(1→ and →4,6)-α-D-Glcp-(1→ may form the patterns of 1→6 linkage. The C1/H6 cross peak at 99.62/3.71 ppm indicates a 1→6 linkage between →4,6)-α-D-Glcp-(1→ and →6)-β-D-Glcp-(1→. The cross peak at 67.78/4.44 ppm suggests a 1→6 linkage between →6)-β-D-Galp-(1→ and →3, 6)-β-D-Galp-(1→.The connection modes of other glycosidic bonds can be also obtained from the HMBC spectrum. The complete structural identification solely based on current data is unattainable since polysaccharides are complex macromolecular motifs with intricate structures. Based on the methylation and NMR data, the backbone structure of the polysaccharide could be tentatively considered as shown in Fig. 6. The main chain of BHPs-W-S3 is →6)-α-D-Glcp-(1→4,6)-α-D-Glcp-(1→6)-β-D-Galp-(1→3,6)-β-D-Galp-(1→, and the major branched chains include β-D-Galp-(1→4)-β-D-Galp-(1→ at O-3 as well as β-D-Glcp-(1→4)-β-D-Glcp-(1→ and β-D-Glcp-(1→4)-β-D-Xylp-(1→ at O-4.

**Fig.6.**
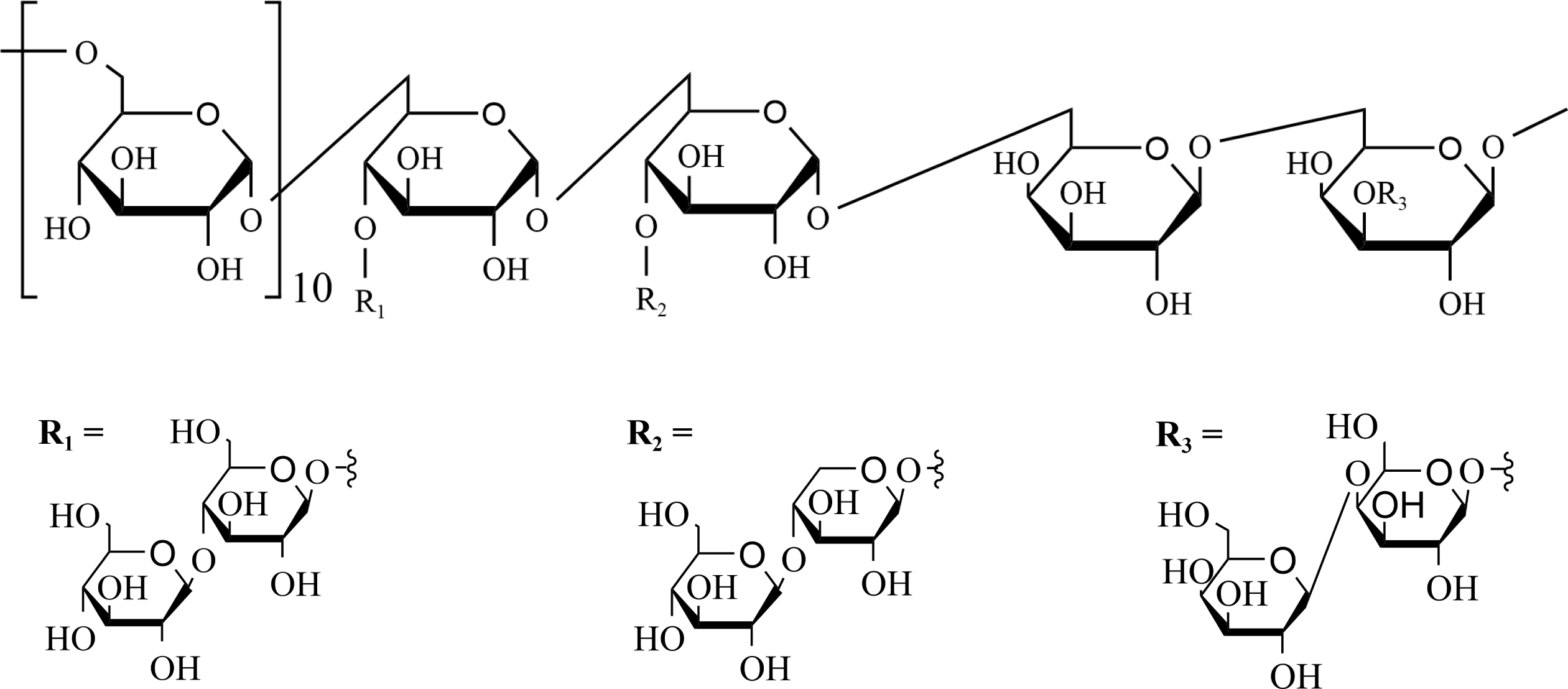
Structural interpolation of BHPs-W-S3.

#### 3.4. Analysis of the triple-helical structure

Congo red can form complexes with the triple-helix β-(1→3)-D-glucan in dilute alkaline solution, which causes a red shift of the maximum absorption wavelength λ_max_ (Guo et al., 2021). Fig. 4C illustrates how the λ_max_ of the Congo red/BHPs-W-S3 complex changes as the NaOH concentration of the solution varies from 0 to 0.8 M. There is a clear rise of λ_max_ when the NaOH concentration increases from 0 to 0.1 M, but λ_max_ varies very little when the NaOH concentration is higher. With Congo red as the control, the Congo red/BHPs-W-S3 complex has stronger absorption over the entire range of 0‒0.8 M NaOH, but there is no significant red shift of the λ_max_. Hence, it can be inferred that BHPs-W-S3 lacks a triple-helical structure.

#### 3.5. *In vitro* anti-inflammatory activity

##### 3.5.1. MTT assay

The MTT assay results show that crude BHPs and BHPs-W-S3 do not have apparent cytotoxic effects on LPS-induced RAW 264.7 cells (Fig. 7A). The results are in line with the findings for lignified okra polysaccharides (Liu et al., 2021). The subsequent studies of anti-inflammatory evaluations thus adopt a concentration range of 25–100 μg/mL for crude BHPs and BHPs-W-S3.

**Fig.7.**
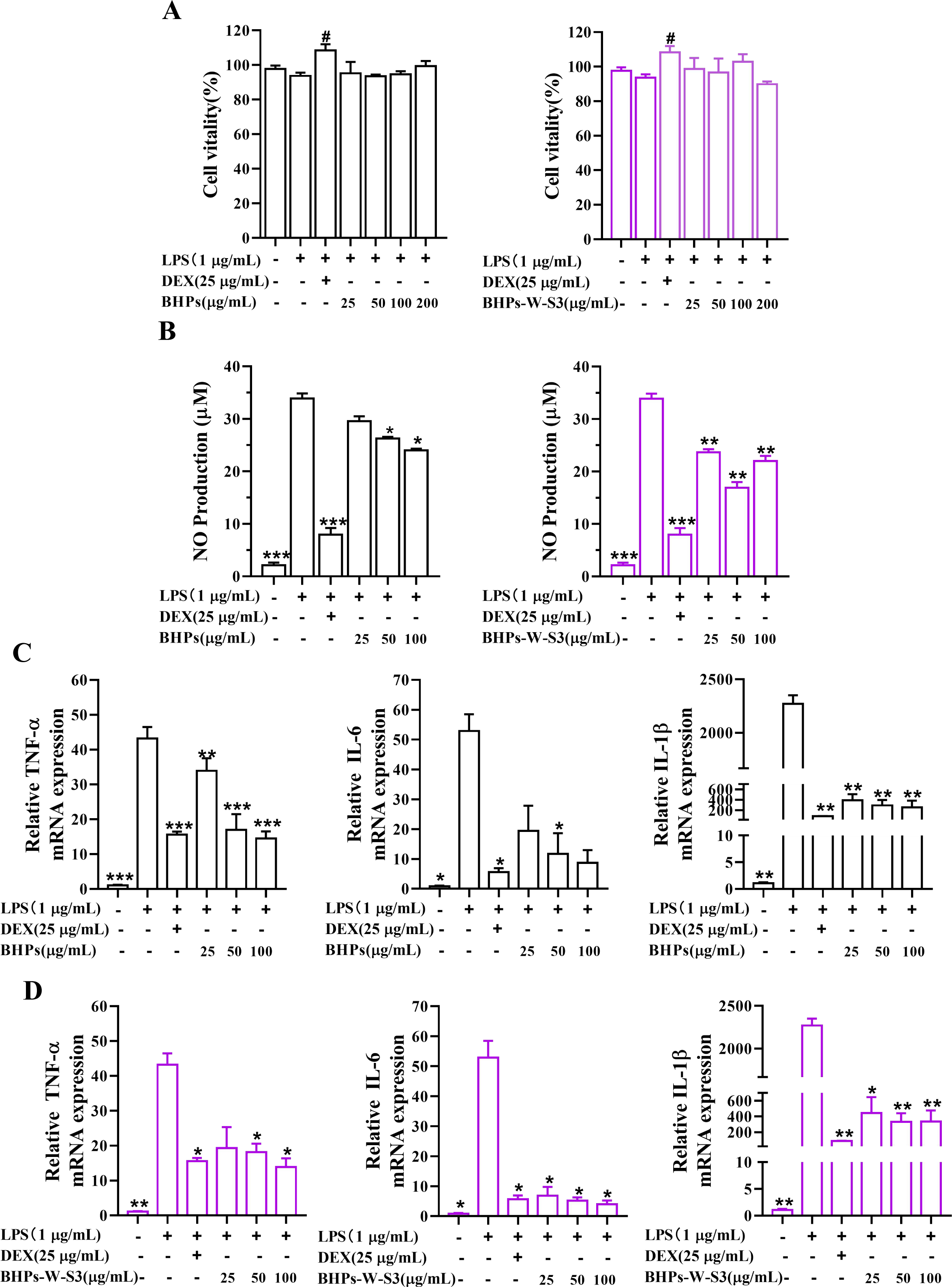
Cell viability and anti-inflammatory effects of crude BHPs and BHPs-W-S3 on RAW 264.7 cells. (A) MTT assay. (B) Griess assay. (C,D) Relative mRNA expressions of TNF-α, IL-6, and IL-1β. All values are presented as means ± SD (n = 3). ^#^ *P <* 0.05, vs. the control group. * *P* < 0.05; ** *P* < 0.01; *** *P* < 0.001, vs. the LPS-induced group.

##### 3.5.2. Anti-inflammatory activity

Activated macrophages can release inflammatory mediators such as NO, prostaglandins, and cytokines (TNF-α, IL-6, IL-1β, etc.) in response to external stimuli (Zhu et al., 2016). Among them, NO plays a pivotal role in a myriad of physiological functions and is intricately linked to inflammatory responses (Surh et al., 2001). When the immune cells are attacked by endotoxins and inflammatory mediators, they can produce NO to initiate a series of immune responses. However, excessive NO generation worsens the inflammatory reaction (Wang et al., 2020), and the NO level can serve as a direct indicator of anti-inflammatory activity.

Nuclear factor kappa-B (NFκB) is an essential intracellular transcription factor involved in immunological responses as well as in the cellular metabolism of several genes (Hu et al., 2018). The stimulation pathway of NFκB primarily involves the breakdown of the phosphorylated IκBα protein to alleviate the inhibition of NFκB. When macrophages are stimulated with LPS or other inflammatory agents, the IκBα kinase participates in the phosphorylation of NFκB, which activates NFκB and allows more NFκB to enter the cell’s nucleus (Ren and Chung, 2007).

Fig. 7B shows that the RAW 264.7 macrophages have significantly higher (*P* < 0.001) NO secretion after the LPS-induced inflammation, while the polysaccharides can moderate the NO level in a dose-dependent manner (*P* < 0.05). The NO level is reduced to 16.97 μM after the application of 50 μg/mL BHPs-W-S3, and the crude BHPs seem to have lower anti-inflammatory activity than BHPs-W-S3. Fig. 7C,D shows the expression of pro-inflammatory cytokines TNF-α, IL-6, and IL-1β in the LPS-induced RAW 264.7 cells is readily moderated by the administration of the polysaccharides (*P* < 0.05).

The western blot results (Fig. 8) show that the phosphorylation levels of both p65 and IκBα in the RAW 264.7 cells are significantly escalated (*P* < 0.01) by the LPS induction and effectively mitigated by the administration of the polysaccharides (*P* < 0.05), while there are no obvious changes in the expression of NFκB and IκBα themselves (Fig. 8C-D and G-H). Hence, it appears that crude BHPs and BHPs-W-S3 inhibit the phosphorylation of p65 and IκBα to reduce the mRNA expressions of related pro-inflammatory cytokines.

**Fig.8.**
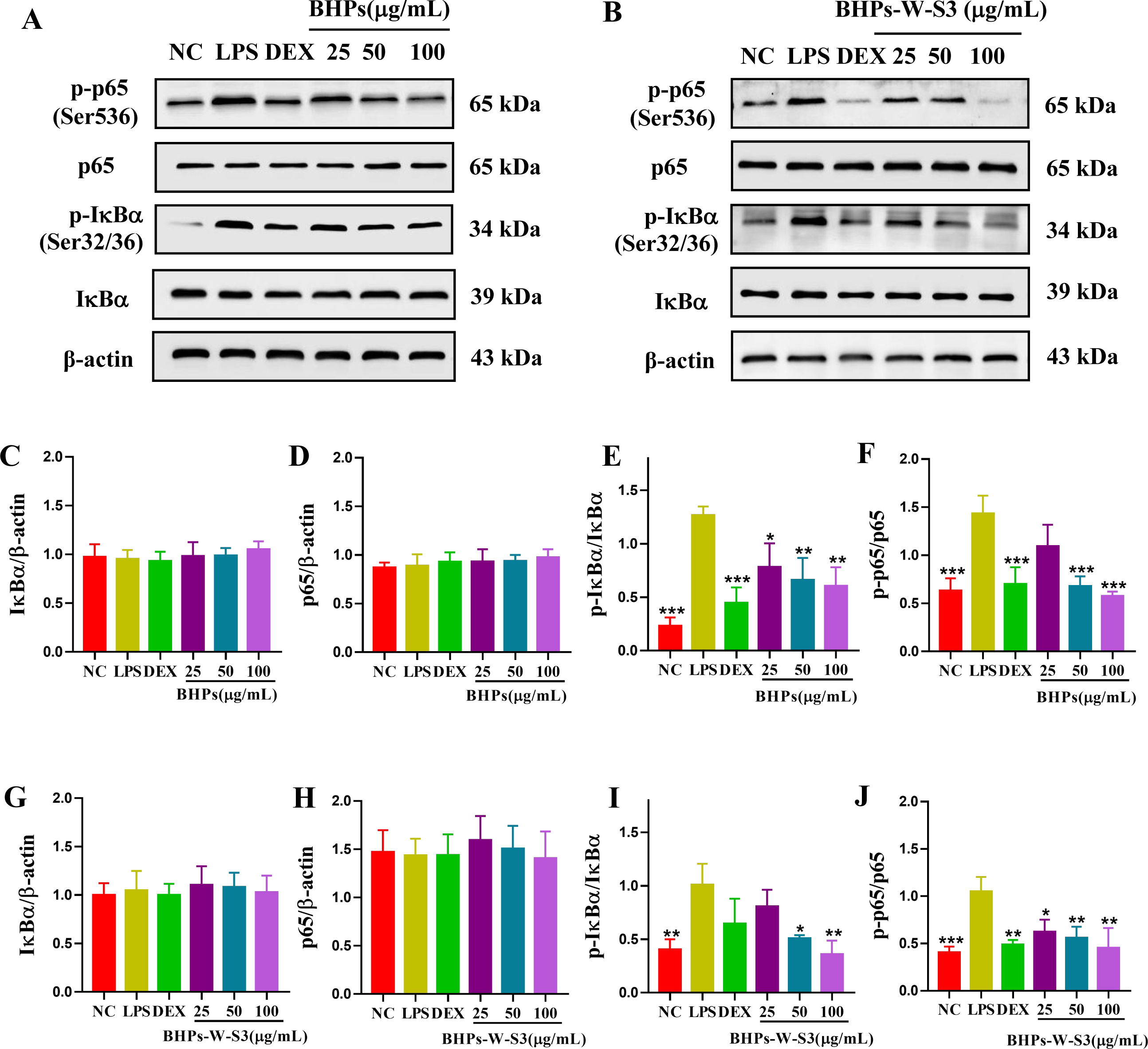
The expression of NFκB pathway-related proteins in RAW 264.7 cells treated with (A) crude BHPs and (B) BHPs-W-S3. Quantification of (C) IκB/β-actin, (D) p65/β-actin, (E) p-IκB/IκB, and (F) p-p65/p65 ratios for RAW 264.7 cells treated with crude BHPs. Quantification of (G) IκB/β-actin, (H) p65/β-actin, (I) p-IκB/IκB, and (J) p-p65/p65 ratios for RAW 264.7 cells treated with BHPs-W-S3. All values are presented as means ± SD (n = 3). * *P* < 0.05; ** *P* < 0.01; *** *P* < 0.001, vs. the LPS-induced group.

##### 3.5.3. Structure-activity relationship

It has been demonstrated that polysaccharides obtained from plants possess anti-inflammatory properties in vitro. The immunomodulatory effects of polysaccharides are closely associated with their physicochemical properties and structural characteristics. The distinct monosaccharide compositions of the polysaccharides, the types of glycosidic linkages, backbone chains and branched structures, molecular weights, and chain conformations all contribute significantly to their biological activities (Dong et al., 2020; Fu et al., 2022; Li et al., 2023). The literature has demonstrated the beneficial impact of glucans on inflammatory conditions (da Nascimento Santos et al., 2014; Li et al., 2017). Similar anti-inflammatory effects of 1, 4, 6-Glcp glycosidic bond have been reported previously (Hu et al., 2021). At present, polysaccharide activity is commonly believed to be greater in the β conformation. Andrea Caroline Ruthes et al. have demonstrated that β-(1→6) and β-(1→3) glycosidic linkages are believed to play a significant role in enhancing anti-inflammatory effects (Ruthes et al., 2013). In the anti-inflammatory assay, *Balanophora* polysaccharides were found to exhibit potent anti-inflammatory activity. Our study reveals that the monosaccharide composition of *BPs* includes glucose, galactose, and mannose, with 1→3 and 1→6 glycosidic linkages. However, it remains unclear whether these structural domains are predominant in the observed anti-inflammatory effects. More research is needed to further understand the structure-activity relationship (SAR) of the polysaccharides from *Balanophora*.

## 3.4. Conclusions

In the current study, extraction procedure of extracting polysaccharides from *B. harlandii* was optimized, and further characterized the detailed structure of the main fraction of polysaccharides BHPs-W-S3. BHPs-W-S3 is a neutral haterpolysaccharided, whose M_w_ is 14.1 kDa. It mainly consists of glucose, galactose, and mannose and has with many linkages, with the main chain having →6)-α-D-Glcp-(1→4,6)-α-D-Glcp-(1→6)-β-D-Galp-(1→3,6)-β-D-Galp-(1→, and the side chains having β-D-Galp-(1→4)-β-D-Galp-(1→ at O-3 along with β-D-Glcp-(1→ 4)-β-D-Glcp-(1→ and β-D-Glcp-(1→ 4)-β-D-Xylp-(1→ at O-4. It lacks a triple-helical structure. The extracted polysaccharides exhibit anti-inflammatory activity when assessed *in vitro* with LPS-induced RAW 264.7 cells. They reduce the release and expression of the pro-inflammatory genes by inhibiting the NFκB pathway. This study provides evidence that BHPs and its main polysaccharide fraction BHPs-W-S3 could be potentially utilized as an anti-inflammatory agent for the prevention and treatment of inflammation-related diseases.

## Declaration of competing interest

The authors declare no conflicts of interest.

### Abbreviations

Ara: arabinose
B. *harlandii*: *Balanophora harlandii* Hook
BHPs: *B. harlandii* polysaccharides
BHPs-W-S3: a neutral polysaccharide of *B. harlandii* polysaccharides
BSA: bovine serum albumin
DEX: Dexamethasone
Fru: Fructose
Fuc: Fucose
FT-IR: Fourier transform infrared spectroscopy
GalA: Galacturonic acid
GC-MS: gas chromatography-mass spectrometry
GlcN: Gal galactose
GlcN: Glucosamine hydrochloride
GlcA: Glucuronic acid
GlcNC: N-Acetyl-D glucosamine
Glc: glucose
GulA: Gullochuronic acid
GPC: gel permeation chromatography
IκBα: inhibitor of kappa Bα
LPS: lipopolysaccharide
ManA: Mannuronic acid
Man: mannose
NMR: nuclear magnetic resonance
NFκB: Nuclear factor kappa-B
PI: polydispersity index
PVDF: polyvinylidene difluoride
PBS: phosphate-buffered saline
p-IκBα: phospho-inhibitor of kappa Bα
Rha: Rhamnose
Rib: Ribose
SD: standard deviation
SAR: structure-activity relationship
TBS-Tween: Tris-buffered saline-Tween
TFA: trifluoroacetic acid
TBS-Tween: Tris-buffered saline-Tween
UV: ultraviolet
Xyl: Xylose

## Acknowledgement

The work was supported by the grants from National Natural Science Foundation of China (Grant No. 81974528 to C.F. Yuan, No. 82174035 to C.F. Yuan, No. 82374107 to C.F. Yuan and No. 81773959 to C.F. Yuan), the innovational group project of Hubei Province Natural Science Foundation in China(Grant No. 2021CFA015 to C.F. Yuan), and the Central Funds Guiding the Local Science and Technology Development (Grant No. 2020ZYYD016 to C.F. Yuan).

## Author contribution

**Yuanyang Li**: Data curation, Formal analysis, Writing–original draft.

**Xueqing Li**: Data curation, Investigation, Writing-review & editing.

**Qi Yuan**: Validation, Resources.

**Leiqi Zhu**, **Fangqi Xia**, **Yaqi Wang**, **Mengzhen Xue**: Formal analysis, Investigation,

**Yumin He**: Conceptualization, Methodology, Writing-Review & Editing, Supervision.

**Chengfu Yuan**: Conceptualization, Methodology, Writing-Review & Editing, Supervision, Project administration, Funding acquisition.

---

**Fig. S1.**
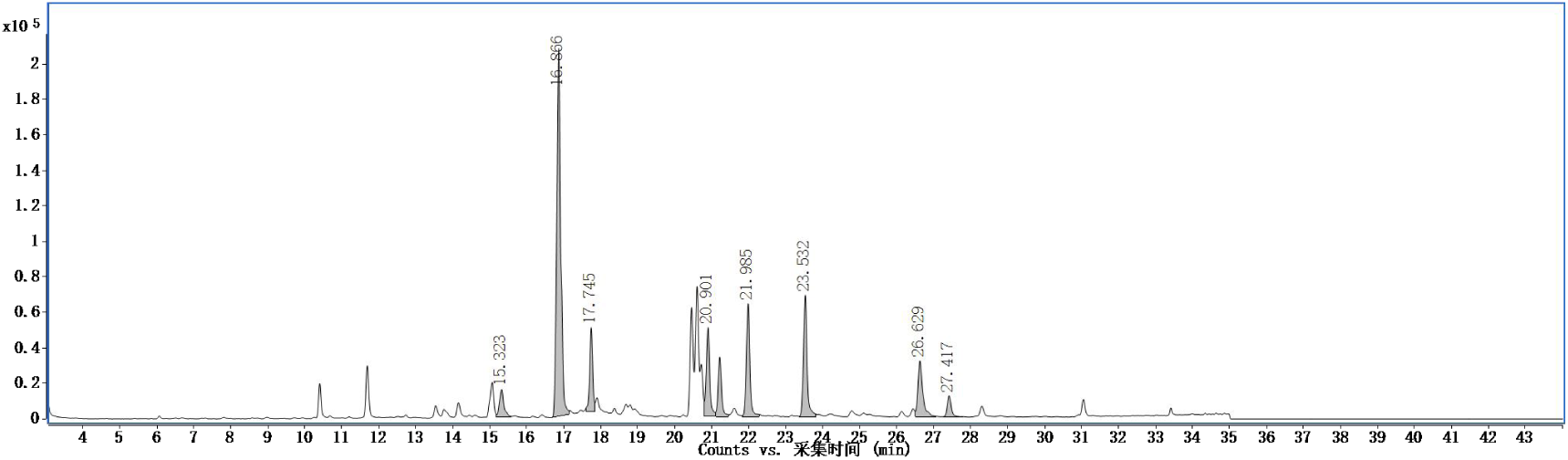
GC-MS chromatogram of methylated BHPs-W-S3.

**Fig. S2.**
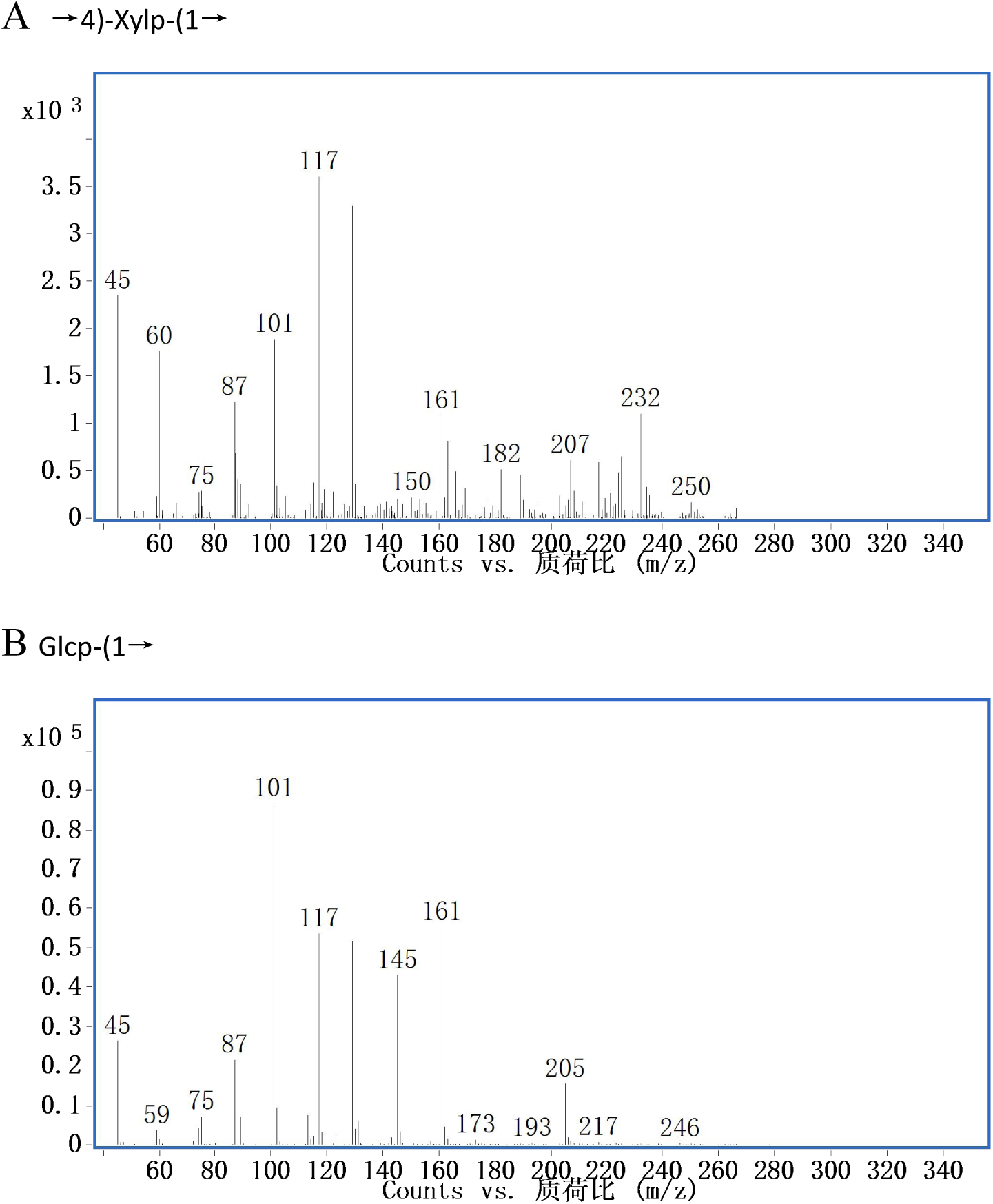
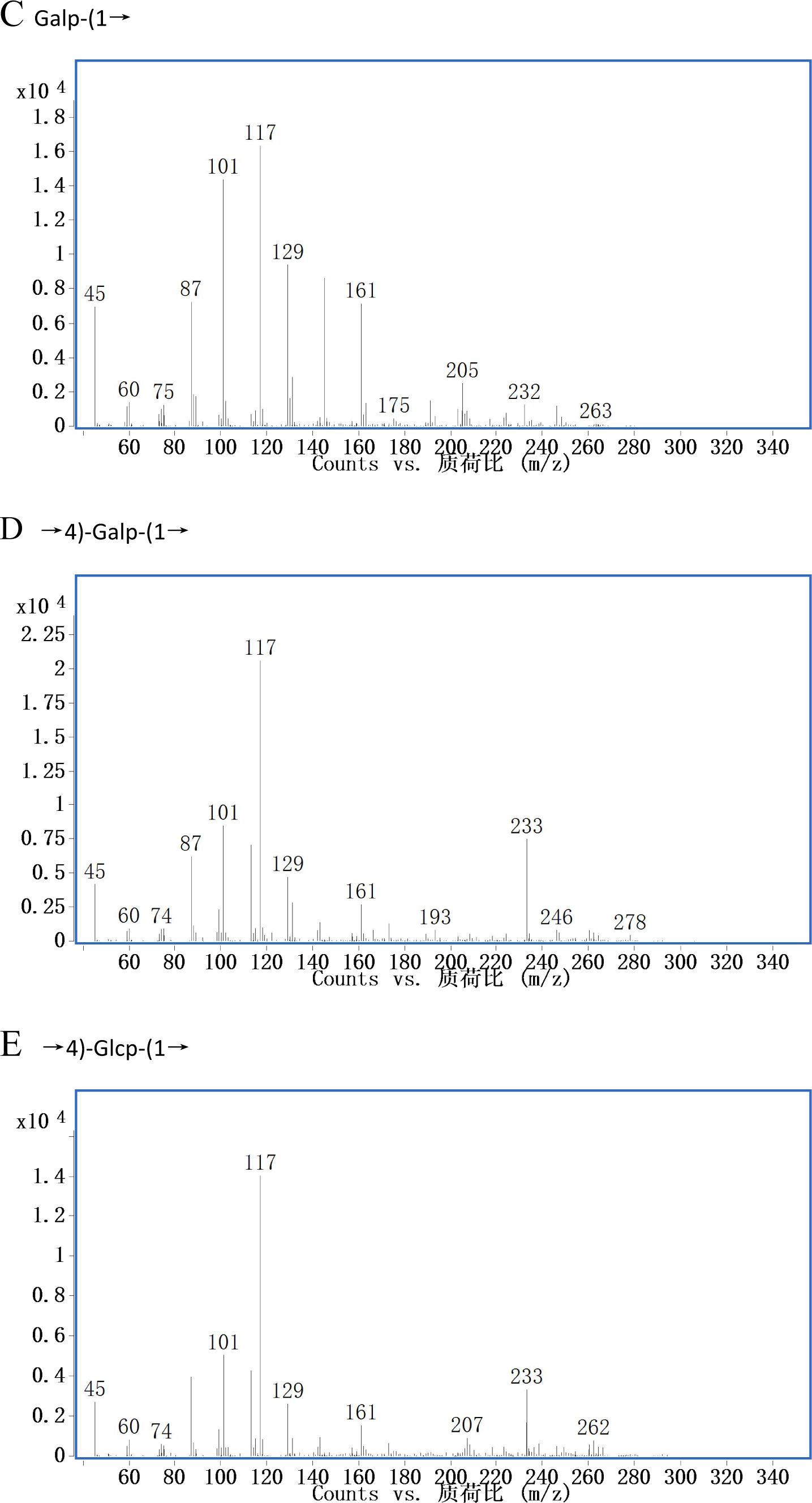
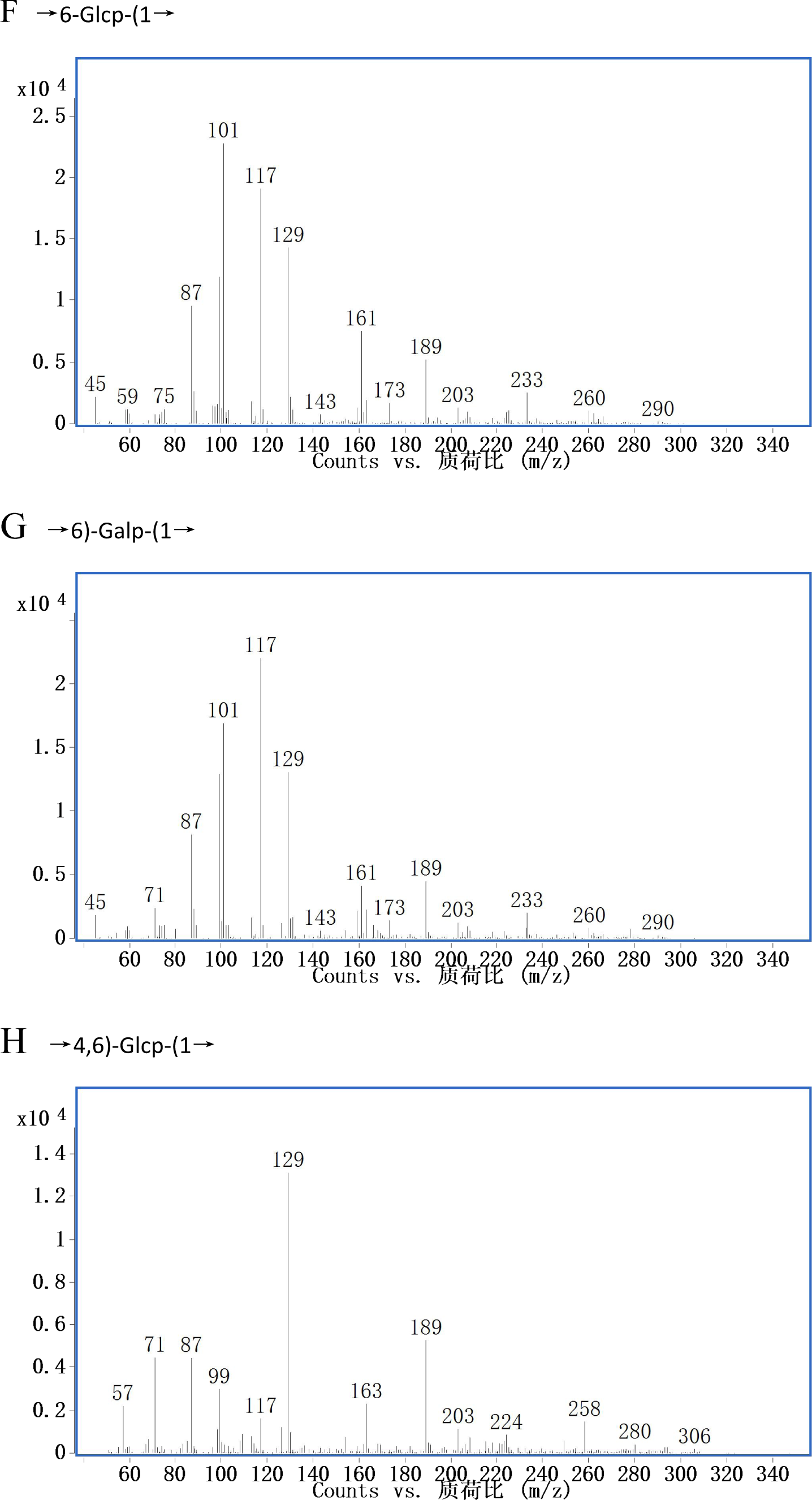
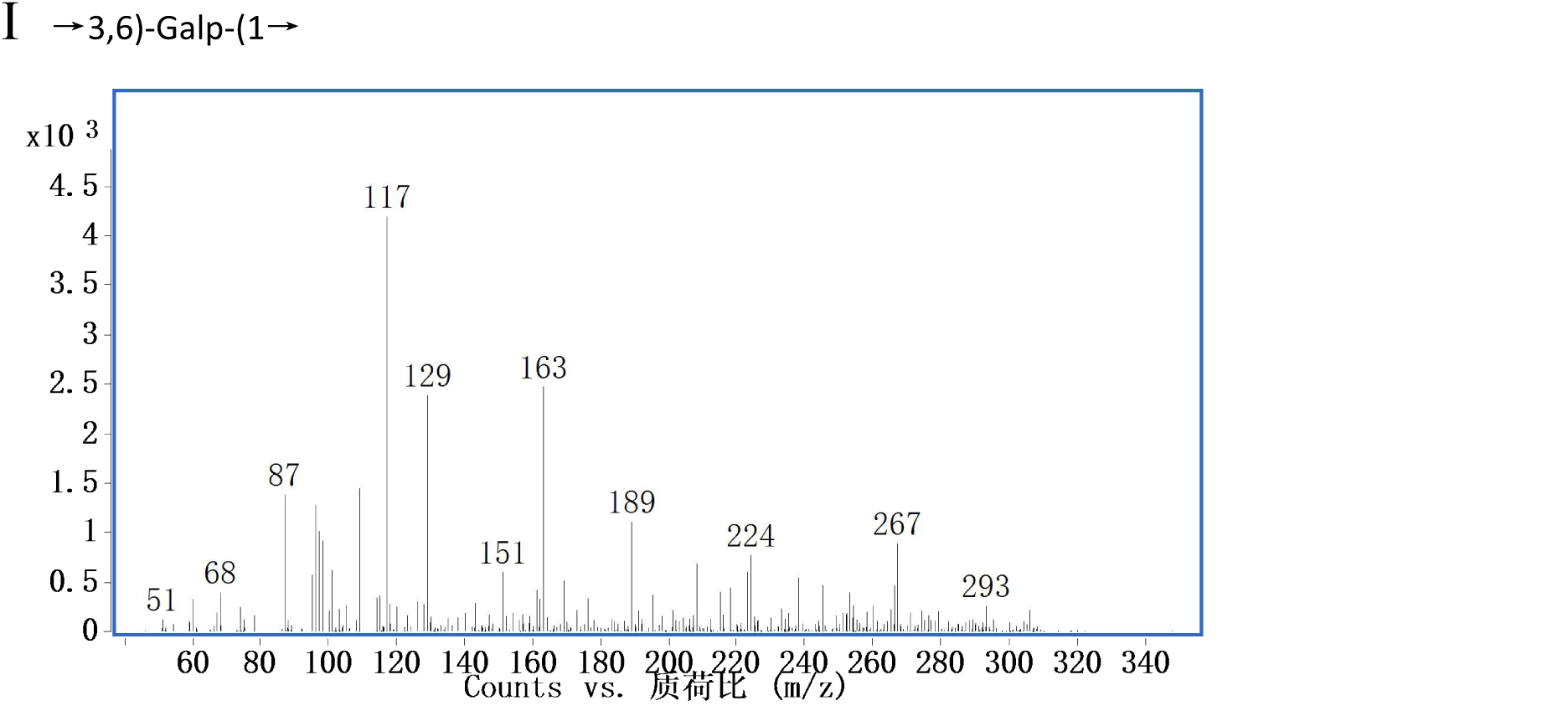
GC-MS fragmentions of the peaks shown in Fig. S1.

**Fig. S3.**
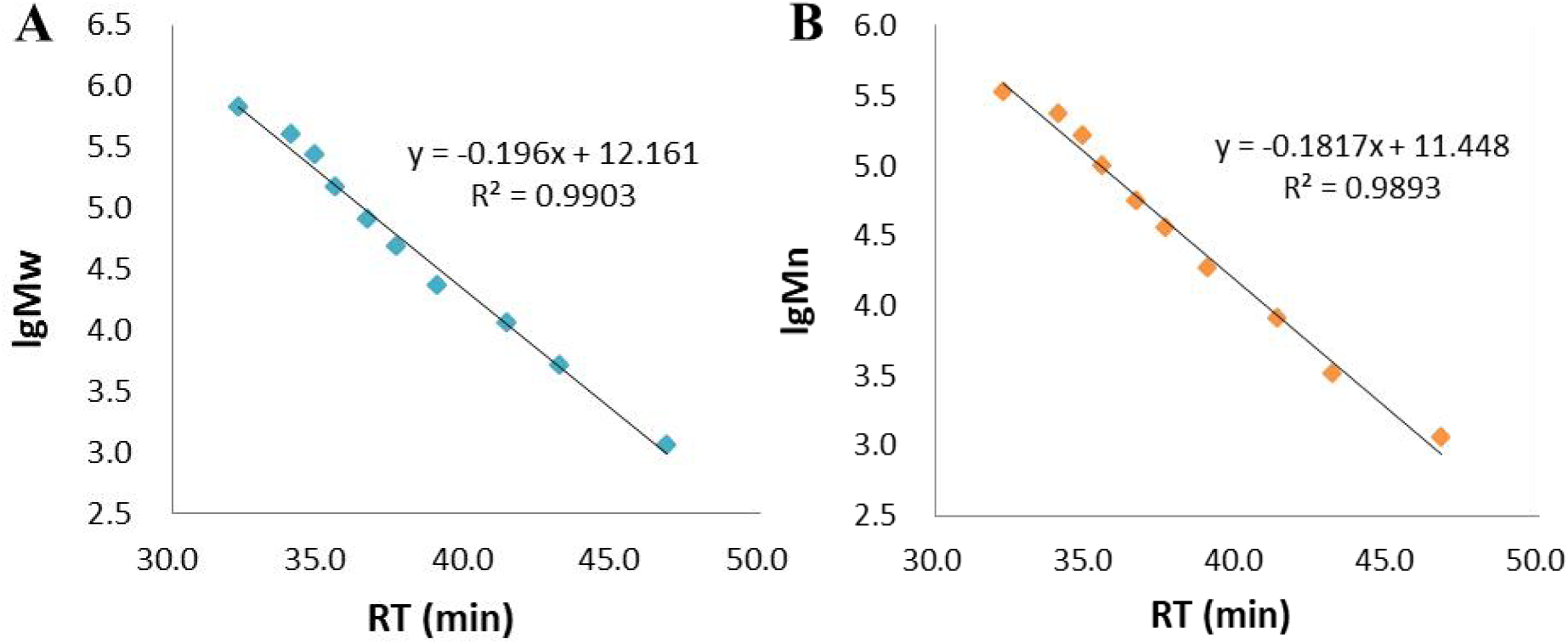
The lgMw-RT(A) and lgMn-RT(B) calibration curve

## References

1. Arab K, Ghanbarzadeh B, Ayaseh A, Jahanbin K (2021) Extraction, purification, physicochemical properties and antioxidant activity of a new polysaccharide from Ocimum album L. seed. International Journal of Biological Macromolecules 180: 643–653

2. Bradford MM (1976) A rapid and sensitive method for the quantitation of microgram quantities of protein utilizing the principle of protein-dye binding. Analytical biochemistry 72: 248–254

3. da Nascimento Santos MS, de Magalhães JEM, Will Castro LSEP, de Sousa Pinheiro T, Sabry DA, Nobre LTDB, Lima JPMS, Baseia IG, Leite EL (2014) Effect of glucans from Caripia montagnei mushroom on TNBS-induced colitis. International journal of molecular sciences 15: 2368–2385

4. Dong Z, Zhang M, Li H, Zhan Q, Lai F, Wu H (2020) Structural characterization and immunomodulatory activity of a novel polysaccharide from Pueraria lobata (Willd.) Ohwi root. International journal of biological macromolecules 154: 1556–1564

5. DuBois M, Gilles KA, Hamilton JK, Rebers Pt, Smith F (1956) Colorimetric method for determination of sugars and related substances. Analytical chemistry 28: 350–356

6. Fu Y-P, Li C-Y, Peng X, Zou Y-F, Rise F, Paulsen BS, Wangensteen H, Inngjerdingen KT (2022) Polysaccharides from Aconitum carmichaelii leaves: Structure, immunomodulatory and anti-inflammatory activities. Carbohydrate Polymers 291: 119655

7. Gu J, Zhang H, Zhang J, Wen C, Zhou J, Yao H, He Y, Ma H, Duan Y (2020) Optimization, characterization, rheological study and immune activities of polysaccharide from Sagittaria sagittifolia L. Carbohydrate polymers 246: 116595

8. Guo C, He Y, Gai L, Qu J, Shi Y, Xu W, Cai Y, Wang B, Zhang J, Zhao Z (2020) Balanophora polyandra Griff. prevents dextran sulfate sodium-induced murine experimental colitis via the regulation of NF-κB and NLRP3 inflammasome. Food & function 11: 6104–6114

9. Guo X, Kang J, Xu Z, Guo Q, Zhang L, Ning H, Cui SW (2021) Triple-helix polysaccharides: Formation mechanisms and analytical methods. Carbohydrate Polymers 262: 117962

10. Herbals TECOC (1999) State administration of traditional chinese medicine.

11. Hu D, Su F, Yang G, Wang J, Zhang Y (2021) Purification, Structural Characterization, and Anti-Inflammatory Effects of a Novel Polysaccharide Isolated from Orostachys fimbriata. Molecules 26: 7116

12. Hu H, Liang H, Wu Y (2015) Isolation, purification and structural characterization of polysaccharide from Acanthopanax brachypus. Carbohydrate polymers 127: 94–100

13. Hu T, Lin Q, Guo T, Yang T, Zhou W, Deng X, Yan J-K, Luo Y, Ju M, Luo F (2018) Polysaccharide isolated from Phellinus linteus mycelia exerts anti-inflammatory effects via MAPK and PPAR signaling pathways. Carbohydrate Polymers 200: 487–497

14. Huang H, Huang G (2020) Extraction, separation, modification, structural characterization, and antioxidant activity of plant polysaccharides. Chemical biology & drug design 96: 1209–1222

15. Jang G, Lee SA, Hong JH, Park B-R, Kim DK, Kim CS (2022) Chondroprotective Effects of 4, 5-Dicaffeoylquinic Acid in Osteoarthritis through NF-κB Signaling Inhibition. Antioxidants 11: 487

16. Jia H, Zhao B, Zhang F, Santhanam RK, Wang X, Lu J (2021) Extraction, structural characterization, and anti-hepatocellular carcinoma activity of polysaccharides from panax ginseng Meyer. Frontiers in Oncology: 4905

17. Kerboua KA, Benosmane L, Namoune S, Ouled-Diaf K, Ghaliaoui N, Bendjeddou D (2021) Anti-inflammatory and antioxidant activity of the hot water-soluble polysaccharides from Anacyclus pyrethrum (L.) Lag. roots. Journal of Ethnopharmacology 281: 114491

18. Kim M, Kim S-R, Park J, Mun S-H, Kwak M, Ko H-J, Baek S-H (2022) Structure and antiviral activity of a pectic polysaccharide from the root of Sanguisorba officinalis against enterovirus 71 in vitro/vivo. Carbohydrate Polymers 281: 119057

19. Li B, Zhang N, Feng Q, Li H, Wang D, Ma L, Liu S, Chen C, Wu W, Jiao L (2019) The core structure characterization and of ginseng neutral polysaccharide with the immune-enhancing activity. International journal of biological macromolecules 123: 713–722

20. Li J, Wang L, Yang K, Zhang G, Li S, Gong H, Liu M, Dai X (2023) Structure characteristics of low molecular weight pectic polysaccharide and its anti-aging capability by modulating the intestinal homeostasis. Carbohydrate Polymers 303: 120467

21. Li L, Zhou G, Fu R, He Y, Xiao L, Peng F, Yuan C (2021) Polysaccharides extracted from balanophora polyandra Griff (BPP) ameliorate renal Fibrosis and EMT via inhibiting the Hedgehog pathway. Journal of Cellular and Molecular Medicine 25: 2828–2840

22. Li Q, Feng Y, He W, Wang L, Wang R, Dong L, Wang C (2017) Post-screening characterisation and in vivo evaluation of an anti-inflammatory polysaccharide fraction from Eucommia ulmoides. Carbohydrate polymers 169: 304–314

23. Li Q, Li J, Li H, Xu R, Yuan Y, Cao J (2019) Physicochemical properties and functional bioactivities of different bonding state polysaccharides extracted from tomato fruit. Carbohydrate polymers 219: 181–190

24. Liu G, Ye J, Li W, Zhang J, Wang Q, Zhu X-a, Miao J-y, Huang Y-h, Chen Y-j, Cao Y (2020) Extraction, structural characterization, and immunobiological activity of ABP Ia polysaccharide from Agaricus bisporus. International Journal of Biological Macromolecules 162: 975–984

25. Liu Y, Li S-M (2020) Extraction optimization and antioxidant activity of Phyllanthus urinaria polysaccharides. Food Science and Technology 41: 91–97

26. Liu Y, Ye Y, Hu X, Wang J (2021) Structural characterization and anti-inflammatory activity of a polysaccharide from the lignified okra. Carbohydrate polymers 265: 118081

27. Luo D, Wang Z, Li Z, Yu X-q (2018) Structure of an entangled heteropolysaccharide from Pholidota chinensis Lindl and its antioxidant and anti-cancer properties. International journal of biological macromolecules 112: 921–928

28. Ma Y, Wang Z, Arifeen MZU, Xue Y, Yuan S, Liu C (2022) Structure and bioactivity of polysaccharide from a subseafloor strain of Schizophyllum commune 20R-7-F01. International Journal of Biological Macromolecules 222: 610–619

29. Qu J, He Y, Shi Y, Gai L, Xiao L, Peng F, Li Z, Wang X, Yuan C (2020) Polysaccharides derived from Balanophora polyandra significantly suppressed the proliferation of ovarian cancer cells through P53-mediated pathway. Journal of Cellular and Molecular Medicine 24: 8115–8125

30. Ren J, Chung SH (2007) Anti-inflammatory effect of α-linolenic acid and its mode of action through the inhibition of nitric oxide production and inducible nitric oxide synthase gene expression via NF-κB and mitogen-activated protein kinase pathways. Journal of agricultural and food chemistry 55: 5073–5080

31. Ren Y, Bai Y, Zhang Z, Cai W, Del Rio Flores A (2019) The preparation and structure analysis methods of natural polysaccharides of plants and fungi: A review of recent development. Molecules 24: 3122

32. Ruthes AC, Carbonero ER, Córdova MM, Baggio CH, Sassaki GL, Gorin PAJ, Santos ARS, Iacomini M (2013) Fucomannogalactan and glucan from mushroom Amanita muscaria: Structure and inflammatory pain inhibition. Carbohydrate polymers 98: 761–769

33. Singleton VL, Rossi JA (1965) Colorimetry of total phenolics with phosphomolybdic-phosphotungstic acid reagents. American journal of Enology and Viticulture 16: 144–158

34. Surh Y-J, Chun K-S, Cha H-H, Han SS, Keum Y-S, Park K-K, Lee SS (2001) Molecular mechanisms underlying chemopreventive activities of anti-inflammatory phytochemicals: down-regulation of COX-2 and iNOS through suppression of NF-κB activation. Mutation Research/Fundamental and Molecular Mechanisms of Mutagenesis 480: 243–268

35. Wang H, Song Z, Xing H, Shi Z, Wu P, Zhang J, Tuerhong M, Xu J, Guo Y (2020) Nitric oxide inhibitory iridoids as potential anti-inflammatory agents from Valeriana jatamansi. Bioorganic Chemistry 101: 103974

36. Wang X, Liu Z, Qiao W, Cheng R, Liu B, She G (2012) Phytochemicals and biological studies of plants from the genus Balanophora. Chemistry Central Journal 6: 1–9

37. Wang Z, Liu X, Bao Y, Wang X, Zhai J, Zhan X, Zhang H (2021) Characterization and anti-inflammation of a polysaccharide produced by Chaetomium globosum CGMCC 6882 on LPS-induced RAW 264.7 cells. Carbohydrate Polymers 251: 117129

38. Xu H-S, Wu Y-W, Xu S-F, Sun H-X, Chen F-Y, Yao L (2009) Antitumor and immunomodulatory activity of polysaccharides from the roots of Actinidia eriantha. Journal of ethnopharmacology 125: 310–317

39. Yuanbei Z, Dadu L, Yunxia LI, Rongdi LI, Lifang W, Lijun Z, Jiahua WU, Shengyuan Z (2017) Research Progress of Balanophora harlandii. Chinese Journal of Ethnomedicine and Ethnopharmacy

40. Zeng X, Li P, Chen X, Kang Y, Xie Y, Li X, Xie T, Zhang Y (2019) Effects of deproteinization methods on primary structure and antioxidant activity of Ganoderma lucidum polysaccharides. International journal of biological macromolecules 126: 867–876

41. Zhang H, Nie S, Cui SW, Xu M, Ding H, Xie M (2017) Characterization of a bioactive polysaccharide from Ganoderma atrum: Re-elucidation of the fine structure. Carbohydrate polymers 158: 58–67

42. Zhu T, Zhang W, Feng S-j, Yu H-p (2016) Emodin suppresses LPS-induced inflammation in RAW264. 7 cells through a PPARγ-dependent pathway. International immunopharmacology 34: 16-24

